# Intracellular amino acid scarcity sensing tunes protein hunger

**DOI:** 10.64898/2026.06.20.732459

**Authors:** Sophie Waldron, Yixin Liu, Vincent Croset

## Abstract

Protein is a key macronutrient that strongly governs food intake. Following dietary protein scarcity, animals display compensatory preference for amino acid-rich food, yet how the brain detects and encodes protein hunger remains unclear. To address this question, we used *Drosophila melanogaster* to investigate the function of peptidergic neurons, a major neuromodulatory cell population that includes key regulators of feeding behaviour and protein homeostasis. We present a high-resolution transcriptional atlas obtained by RNA sequencing of single peptidergic nuclei isolated from the brains of sated and amino acid deprived flies. Consistent with a resource-sparing response, differential expression analysis revealed widespread transcriptional downregulation in the amino acid deprived state. A screen of differently expressed candidates identified *Lsp2*, a gene encoding an amino acid storage protein, as a regulator of protein intake. Knock down and overexpression experiments demonstrate that Lsp2 levels act as a gauge in a specific population of peptidergic neurons expressing *Myoinhibiting peptide precursor* (*Mip*) to modulate protein appetite. We further show that compensatory protein feeding following amino acid deprivation depends on maintaining *Mip*+ neuron activity within an appropriate range. Our findings reveal that peptidergic neurons can directly sense intracellular protein deficiency and use this information to guide protein intake. By linking amino acid storage to the neuronal control of feeding, this work establishes a foundation for understanding how protein needs are encoded in the brain.

## Introduction

Beyond the general urge to consume food under starvation, hunger also manifests as specific appetites, guiding organisms to consume foods rich in dietary nutrients they are lacking. Protein is a dietary macronutrient with a strong homeostatic drive^1,2^, which makes a major contribution in determining total energy intake^3–6^. Most animals display compensatory preference for amino acid rich food following dietary scarcity^2,7,8^, even when protein deprivation is slight^9,10^ or limited to essential amino acids^11–14^.

Protein homeostasis is regulated at multiple levels, from peripheral sensory systems^15^, internal amino acid sensing in the gut^16^, and neural circuits in the CNS^17^. In amino acid deprived mammals, chemosensory information processed in the anterior piriform cortex leads to rapid rejection of a food source lacking essential amino acids^18,19^. In *Drosophila*, gustatory receptor neurons in the mouthparts, legs and wings that respond to a fly’s main source of protein, yeast, increase their response to these food stimuli following protein deprivation^2,15,20^. Inhibiting these peripheral responses suppresses yeast preference under deprivation conditions^2^.

Across species, intracellular sensing of amino acid and ATP levels is achieved through the TOR/S6K pathway^21,22^, which controls the propensity to consume protein-rich food sources following mating or protein deprivation^23^. Yet, the neuronal circuits controlled by these signals have not been specified.

Peptidergic systems play an essential role in feeding and appetite regulation^24,25^. Secreted peptides are also involved in sucrose^26,27^ and protein homeostasis, particularly by mediating the satiating and consumption-suppressing effects of protein. In humans and other mammals, anorexigenic gut peptides such as peptide YY, peptide-1 and glucagon^28^, as well as neuropeptide Y expression in the hypothalamus^9^, signal for protein satiety. In *Drosophila,* fat body-secreted FIT acts as a protein satiety signal on insulin producing cells (IPCs), which secrete Ilp2 to regulate the cessation of food consumption commensurate with protein intake^29^.

Less is known about neuronal circuits involved in compensatory feeding following dietary protein depletion. In Drosophila, protein consumption following deprivation is driven by a pair of PPM2 dopaminergic “wedge” neurons, which evoke persistent protein seeking and decrease carbohydrate preference^17^.

Given the role of neuropeptides as major determinants of feeding behaviour, we sought to identify molecular and cellular underpinnings of protein hunger control by peptidergic systems. Using single-nucleus RNA-sequencing, we identified broad and cell type-specific transcriptional changes triggered by protein hunger across peptidergic cell populations, which lead to the discovery that certain peptidergic neurons leverage the depletion of an amino acid storage protein to sense amino acid scarcity and drive protein consumption.

## Results

### Peptidergic signalling drives intake of proteins, but not sugar

We first confirmed that protein deprivation increases yeast consumption under our lab’s conditions by quantifying sugar and yeast consumption in amino acid- or sugar-deprived mated females. Using FlyPAD, a behavioural platform that tracks prandiology across two food sources in real-time^30^, we quantified the number of sips on 10% yeast or 3% sugar. Measurements were taken during the hour following three days of *ad libitum* exposure to chemically defined diets lacking either amino acids or sugar^11^ (Figures 1A and 1B). Amino acid deprived flies consumed almost three times more yeast than those kept on a complete diet or deprived of sugar (Figures 1B and 1D). Consistently, sugar-deprived animals consumed twice more sucrose than the two other groups (Figures 1C and 1D). This was evident across related consumption and food interaction measures including sips (Figures 1 B-D), feeding bursts (sequence of consecutive sips; Figures S1A and S1B) and activity bouts (periods of frequent food interactions; Figures S1C and S1D).

**Figure 1.**
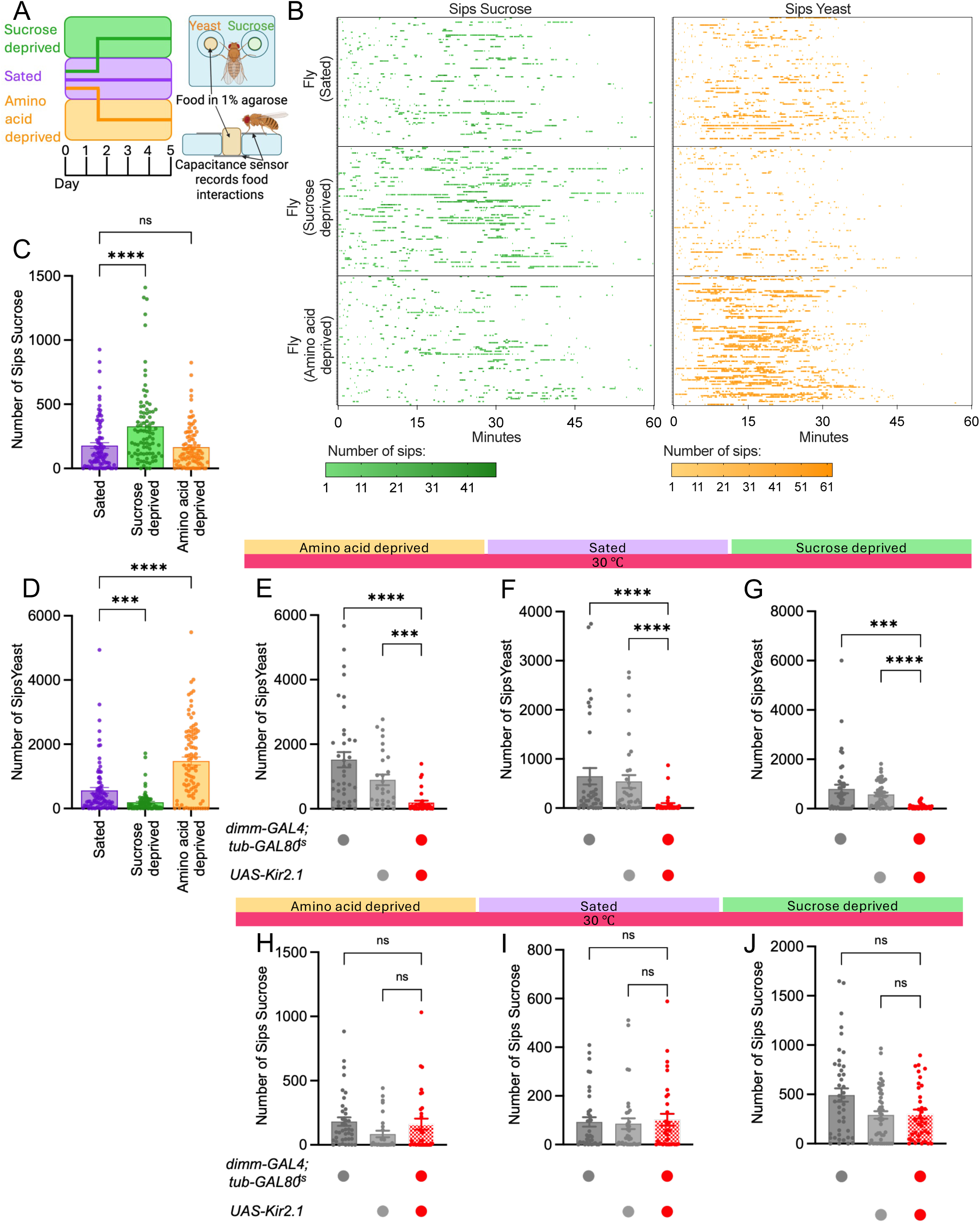
Protein homeostasis is under peptidergic control in amino acid deprived Drosophila. (**A**) Schematic of nutrient deprivation protocol and FlyPAD. (**B**) Feeding raster tracking sips on sugar and yeast for sated, sucrose deprived and amino acid deprived flies. n = 86-90 flies per group. (**C-D**) Barplots representing increase in sucrose consumption after 3 days of sugar deprivation (C) and increase in yeast consumption and decrease in sugar consumption after 3 days of amino acid deprivation (D). n = 86-90 flies per group. (**E-G**) Barplots showing that suppressing excitability in peptidergic neurons with Kir2.1 abolishes yeast consumption in amino acid deprived (E), sated (F), and sugar-deprived (G) flies. n = 28-48 flies per group. (**H-J**) Barplots showing that suppressing excitability in peptidergic neurons with Kir2.1 does not affect sugar consumption in amino acid deprived (H), sated (I), and sugar-deprived (J) flies. n = 28-48 flies per group. ns p ≥ 0.05, *** p < 0.001, **** p < 0.0001, Kruskal-Wallis ANOVA with Dunn’s multiple comparisons test. Barplots are mean +/- standard error of the mean (SEM).

To test whether peptidergic neurons as a population are required to drive yeast consumption, we expressed the Kir2.1 inward-rectifier potassium channel under the *dimm-Gal4* driver, to specifically suppress excitability in these neurons^31–34^. We temporarily restricted Kir2.1 to the final day of the deprivation and test phase, using a ubiquitously expressed temperature-sensitive variant of the Gal4 inhibitor GAL80 (*tub-GAL80ts*) to temporally restrict Kir2.1 expression^35^. Hyperpolarisation of peptidergic neurons drastically suppressed yeast consumption in protein-deprived (Figure 1E), sated (Figure 1F), and sugar-deprived (Figure 1G) animals, demonstrating a state-independent failure in protein homeostasis. Sugar intake was not affected in any of these feeding states (Figures 1H-1J). Genetically identical animals kept at restrictive temperature showed unaltered yeast consumption (Figure S1E-S1G). These results indicate that, as a population, peptidergic neurons promote yeast appetite, irrespective of nutritional states. They also add to the growing body of evidence suggesting that peptidergic hunger signals primarily control the motivation to consume protein, rather than representing a general drive for food consumption^36,37^.

### RNA-sequencing of single peptidergic nuclei

Peptidergic neurons constitute a heterogeneous population including neurons regulating metabolism, stress responses, reproduction, osmoregulation, circadian activity, and nutrition^25,38–43^. To address the transcriptional changes occurring in individual peptidergic neuron types during protein deprivation, we performed single-nucleus sequencing (snRNA-seq) selectively on these neurons isolated from protein-deprived flies and controls on a complete diet. We expressed the GFP-tagged nuclear protein unc84^44^ in *dimm-Gal4* nuclei (Figures 2A and S2A), and used fluorescence activated sorting to isolate GFP+ nuclei from dissociated brains^45^ (Figure S2B). Nuclei were processed with the 10X Genomics Chromium system to generate an atlas of single peptidergic nuclei. We retrieved a total of 15,505 transcriptomes (Figure S2C), with an average of 48,477 reads each. Using supervised clustering, we classified nuclei according to their peptide identity, obtaining a total of 36 clusters (Figures 2B and 2C), containing between 91 and 1166 nuclei each (Figure 2D).

**Figure 2.**
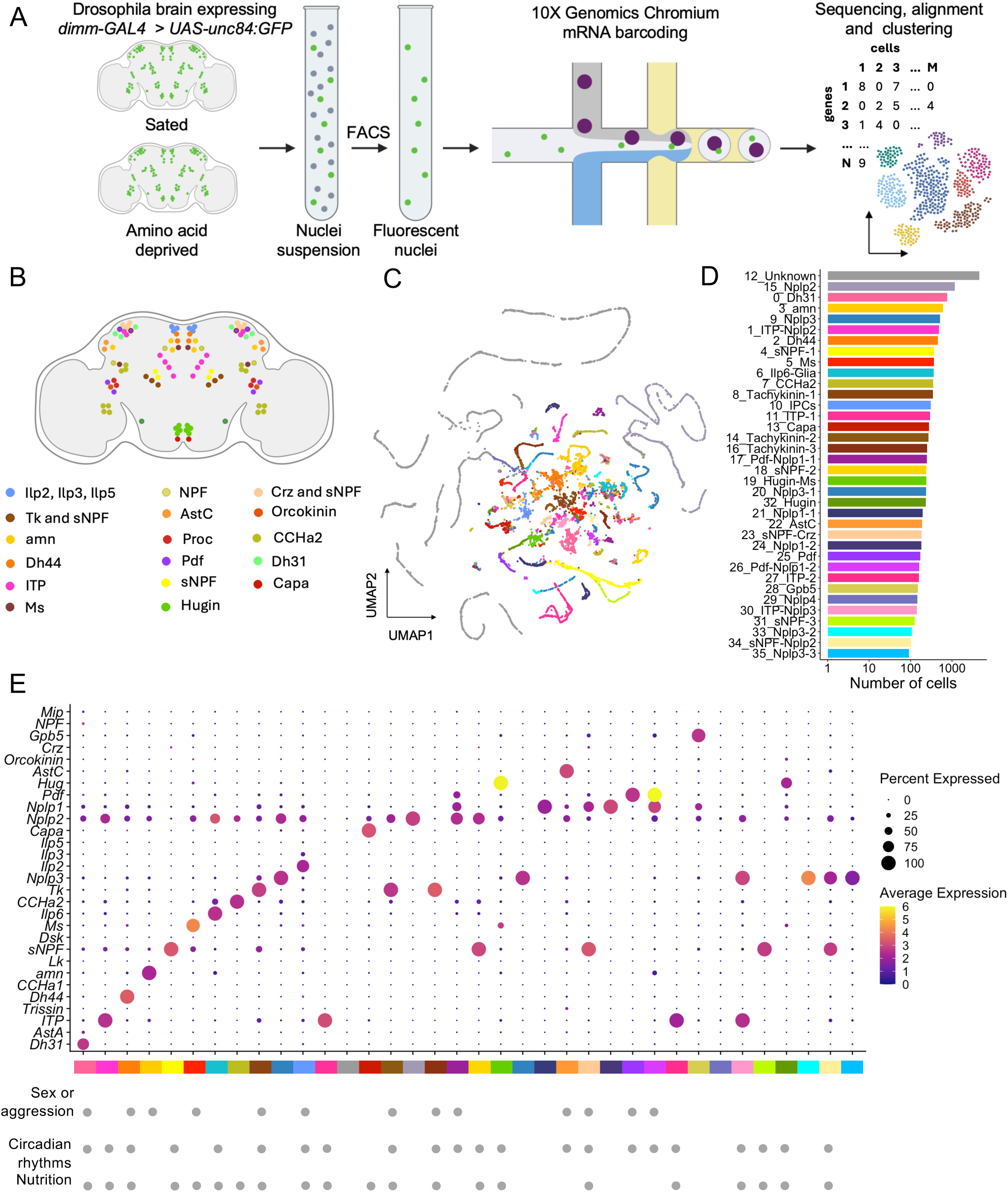
A single-cell atlas of peptidergic cells from the Drosophila brain. (**A**) Single nuclei RNA-sequencing procedure. Nuclei from sated and amino acid deprived flies are sorted to collect GFP-expressing nuclei from peptidergic neurons. RNA from these is then barcoded using 10X Genomics Chromium, before sequencing and further analysis. (**B**) Schematic of the distribution of peptidergic cell types in the *Drosophila* brain. (**C**) UMAP plot representing peptidergic nuclei clusters. (**D**) Number of cells in each cluster, colour coded as in (C). (**E**) Dot plot showing expression of known neuropeptides (Y-axis) across peptidergic clusters (X-axis), colour coded as in (C). Dot intensity represents the average normalised expression level (Log2FC) and diameter represents the fraction of cells expressing the gene per cluster. Grey dots below the graph indicate known roles of peptidergic cell types based on existing literature.

Marker analysis revealed neuropeptide-encoding genes among the top markers of 34 clusters (Figure 2E and Data S1), confirming that these correspond to known peptidergic neuron types. Most of these clusters display characteristic marker combinations consistent with previous studies^46,47^. Interestingly, transcriptionally distinct clusters emerged for some populations previously considered homogeneous. For example, *Hugin*+ and *Pdf*+ neurons separate into distinct clusters defined by differential expression of transcription factors and markers associated with distinct neurotransmitter systems, consistent with possible co-transmission (Figures S2E and S2F and Data S1). One cluster contained glia^45^, and another contaminant nuclei lacking neuropeptide expression, reflected by their elongated, linear arrangement in the UMAP space (Figure 2C)^48,49^. Non-peptidergic clusters were excluded from further analyses. Overall, our annotation supports the biological relevance of the atlas and provides a framework for comparing transcriptional changes among feeding states.

### Amino acid deprivation changes the transcriptional landscape of peptidergic neurons

Changes in gene expression resulting from amino acid deprivation were calculated for each peptidergic cluster individually. We previously demonstrated that correcting for biases induced by zero-inflation increases the accuracy of differential gene expression analysis in single-cell data from full *Drosophila* brains^50^, a step made even more relevant by the higher sparsity of single-nucleus data. We therefore used ZINB-WaVE^51,52^ to model count distributions and estimate data dispersion, prior to quantifying differential expression^53^.

Consistent with the need to conserve resources under conditions of scarcity, we found that most differential expression events involved genes that were downregulated under amino acid deprivation. Twenty genes were overexpressed, spread across 5 clusters, whereas 17 were downregulated (Figure 3A). Inspection of differential expression across clusters (Figure 3B) revealed a marked asymmetry, with most upregulated genes exhibiting cluster-specific regulation while downregulated genes were more broadly regulated, sometimes across a majority of clusters (Figures 3B and 3C).

**Figure 3.**
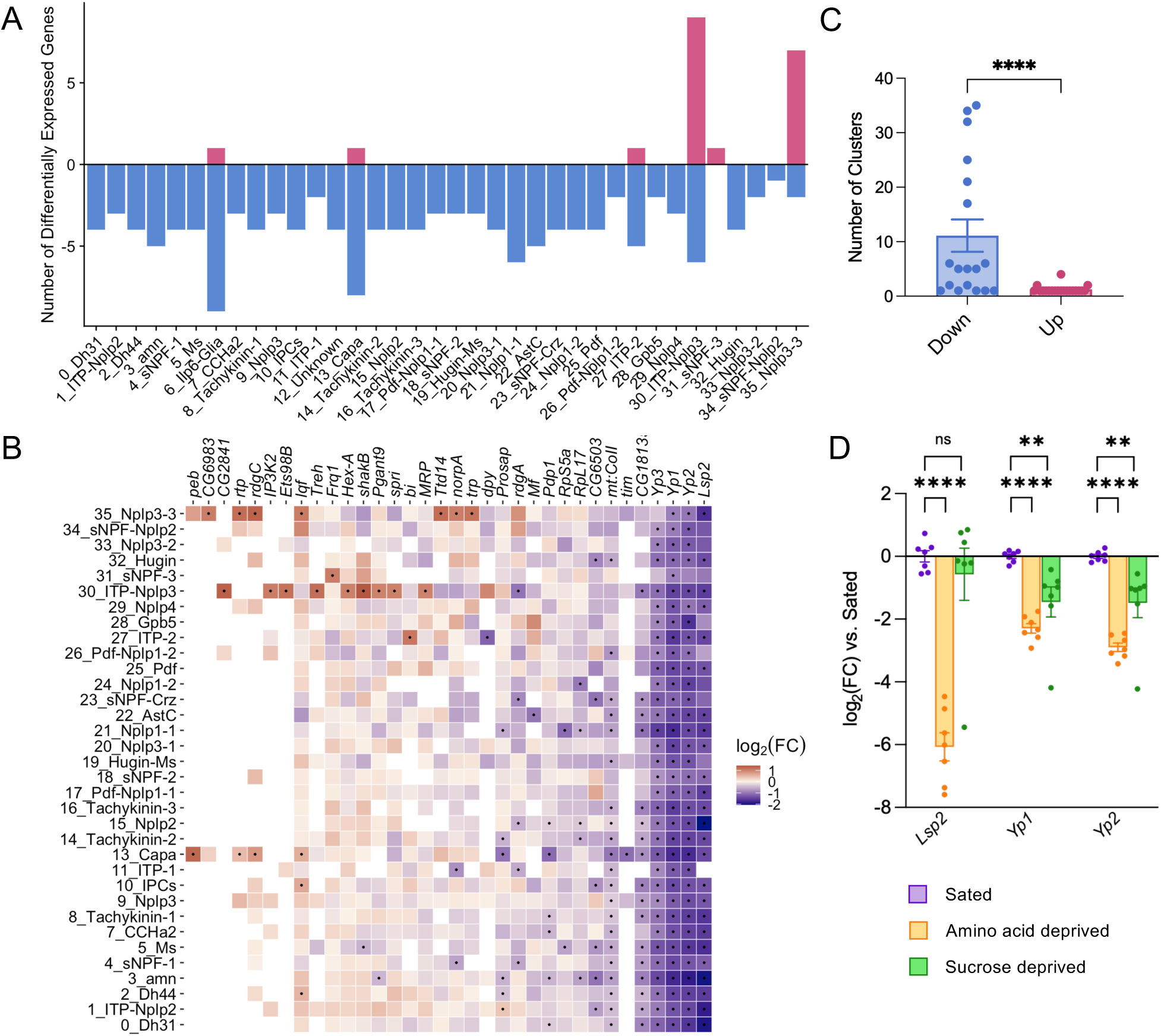
Transcriptional signature of protein deprivation in peptidergic clusters. (**A**) Number of differential expression events (Log_2_(FC) > 1 & adjusted p-value <0.05) across peptidergic cell clusters. Positive values represent upregulated genes while negative values represent downregulated genes. (**B**) Heatmap showing fold change for each significantly regulated gene in amino acid deprived *vs.* sated flies, across all peptidergic clusters. Black dots: adjusted p-value < 0.05 (Ward test with Benjamini-Hochberg correction). Empty tiles: transcripts below detection threshold. (**C**) Number of peptidergic clusters showing significant upregulation or downregulation of each differentially expressed gene (adjusted p-value < 0.05). Downregulated genes are more broadly regulated than upregulated genes. n = 34. (**D**) Relative expression changes measured by RT-qPCR for *Lsp2*, *Yp1* and *Yp2* in the heads of amino acid or sucrose deprived flies, relative to sated flies. n = 6-7 groups of 25 flies each. ns p ≥ 0.05, ** p < 0.01, **** p < 0.0001, Kruskal-Wallis ANOVA with Dunn’s multiple comparisons test. Individual data points are individual genes in (C) and groups of 25 flies in (D). Barplots are mean +/- standard error of the mean (SEM).

Many upregulation events happened in clusters expressing *Nplp3* and/or *ITP*, and involve genes linked to nutrient metabolism and feeding behaviour. *CG6983* encodes a putative adipose secreted signaling protein and its mammalian orthologue is important for glucose homeostasis^54^. The trehalose hydrolysis enzyme *Treh* is essential for haemolymph glucose and water homeostasis^55^, with loss-of-function mutant larvae showing reduced adaptability to low protein diets^56^. *Hex-A*, a sugar metabolism enzyme, acts as a glucose sensor in larval insulin producing cells^57^. Finally *MRP* expression in the malpighian tubes regulates overall food intake^58^.

Surprisingly, despite established links between protein deprivation and inhibition of neuronal TOR^23^, no genes directly associated with TOR activity (GO:0031929) were differently regulated. However, some differently regulated genes had links to the nutrient-sensing TOR/SK6 pathway. *spri*, upregulated in ITP-Nplp3 cluster, is a potential Ras effector^59^, whereas liquid facets (*lqf)*, upregulated in Dh44, IPCs, Capa and Nplp3-3 clusters regulates autophagy downstream of TOR^60^. Downregulation of the diacylglycerol kinase *rdgA* may reduce pS6K to control lifespan and response to oxidative stress^61^.

The yolk proteins *Yp1*, *Yp2*, *Yp3* and larval serum protein 2 *(Lsp2)* are strongly downregulated in most peptidergic clusters. These genes encode amino acid storage proteins, which serve as a nutrient reservoir for the developing embryo^62–64^, and have known interactions *in vitro*^65^. Parallel upregulation of *lqf* may promote endocytosis of these proteins to meet cellular amino acid demands^60^. These amino acid storage proteins are also present in adult *Drosophila* tissues, including the female adult brain^66,67^. In the case of *Lsp2*, a hexamerin, the adult peptide is larger than the larval one, suggesting a possible divergent function^68^. In ovarian follicle cells and adipose tissue, *Lsp2* acts as an effector of mTOR pathway to promote germline stem cell maintenance and lifespan, respectively^69–71^. This indicates a possibility for a signalling role for Lsp2 alongside its function as a nutrient reservoir. Although *Lsp2* downregulation has already been measured in the brain under starvation^72,73^, we confirmed downregulation of *Yp1*, *Yp2* and *Lsp2* under amino acid deprivation using RT-qPCR on heads from amino acid and sucrose deprived flies. While *Yp1* and *Yp2* were downregulated under both protein and sugar deprivation, *Lsp2* was specifically repressed in protein deprived animals (Figure 3D). This points to the possibility of *Lsp2* playing a specific role in the response to protein deprivation.

### Lsp2 is a novel regulator of protein consumption

To identify differently regulated genes involved in the control of protein consumption, we used RNA interference (RNAi) to address the function of a selection of five broadly down-regulated candidates (*Yp1*, *Yp2*, *Yp3*, *rdgA* and *Lsp2*). For each gene, RNAi expression was induced during the three-day protein deprivation and test using *GAL80^ts^*. We hypothesised that downregulating these candidates under sated conditions could mimic protein deprivation and increase yeast consumption; however, this was not the case (Figures S3A-E). Similarly, targeting *Yp1*, *Yp2*, *Yp3*, or *rdgA* in protein-deprived animals did not affect food consumption (Figure 4A-D); however, knocking down *Lsp2* significantly increased yeast consumption (Figure 4E), a phenotype replicated using a second, independent RNAi strain (Figure 4F).

**Figure 4.**
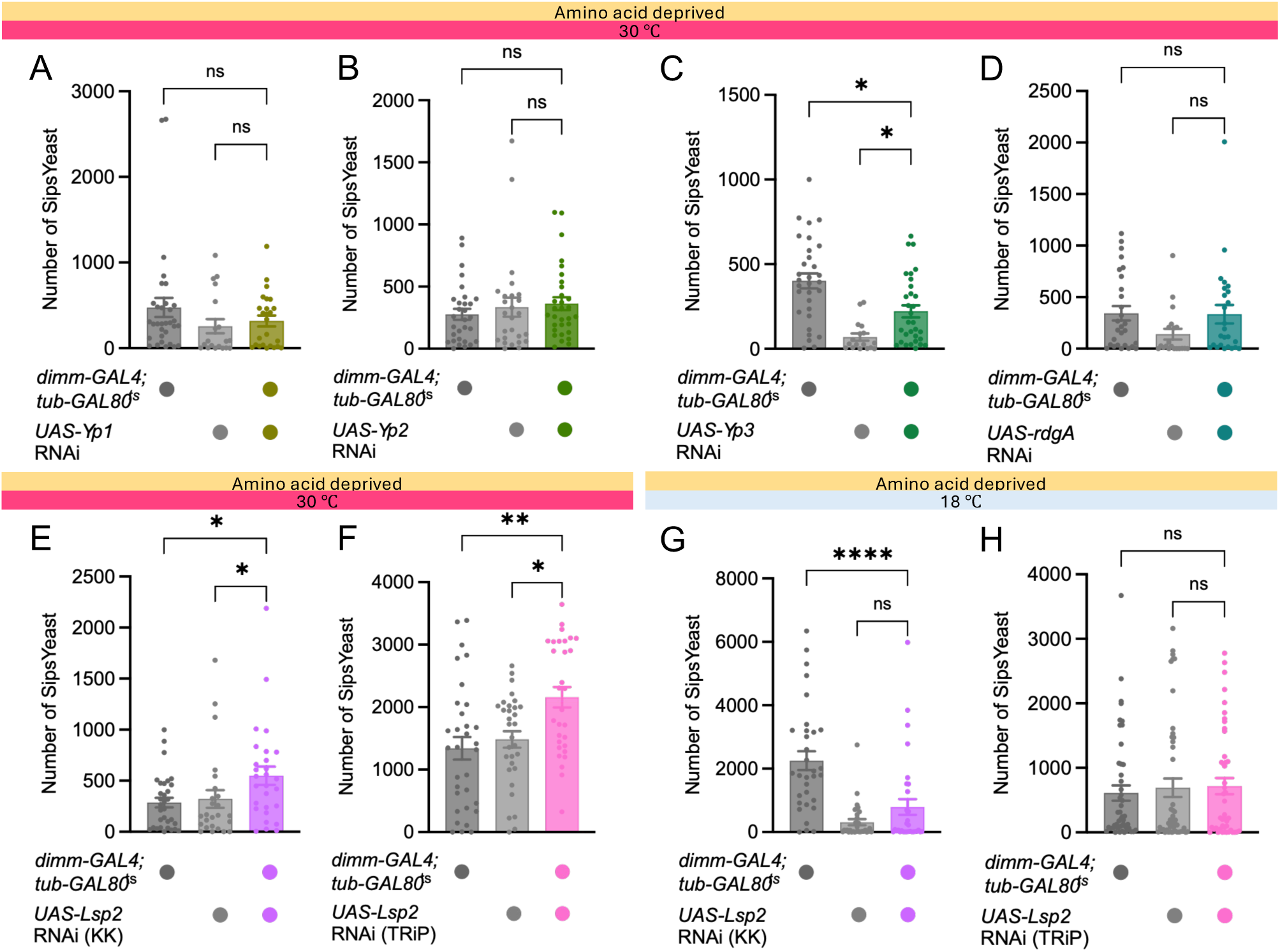
Lsp2 knock down increases protein intake. (**A-D**) Quantification of yeast intake in protein deprived flies expressing RNAi against *Yp1* (A), *Yp2* (B), *Yp3* (C), and *rdgA* (D) in peptidergic neurons using *dimmed-Gal4* driver. No significant effect was observed. n = 17-32 flies per group. (**E-F**) Increased yeast intake was measured in protein deprived flies expressing two independent RNAi transcripts against *Lsp2* in peptidergic neurons using *dimmed-Gal4* driver. n = 25-32 flies per group. (**G-H**) No significant change in yeast consumption was measured in flies genetically identical to those used in (E) and (F) kept at 18°C. n = 32-47 flies per group. ns p ≥ 0.05, * p < 0.05, ** p < 0.01, **** p < 0.0001, Kruskal-Wallis ANOVA with Dunn’s multiple comparisons test. Individual data points are single flies. Barplots are mean +/- standard error of the mean (SEM).

Genetically identical controls kept at permissive temperature throughout the experiment did not display any difference in yeast consumption (Figures 4G and 4H). *Lsp2* knock down did not affect yeast or sugar consumption in sugar-deprived animals (Figure S3F). Together, these findings suggest that reduced *Lsp2* expression in peptidergic neurons is sufficient to promote yeast consumption in protein-deprived flies, whereas this effect is suppressed in sated flies.

### Lsp2 signalling is independent of autophagy and proteolysis

Macroautophagy, the lysosomal degradation of cytoplasm for nutrient recycling, is essential for the mobilisation of amino acids upon starvation^74,75^. This process is controlled by mTOR signaling, which plays a direct role in protein homeostasis^23^, and is modulated by Lsp2 in the fat body^70^. We reasoned that preventing autophagy may result in an increased state of protein deprivation, with flies unable to mobilise amino acids from Lsp2 degradation, and lead to increased yeast consumption. Inhibition of autophagy with temporally controlled expression of RNAi against *Atg5*^76,77^ and *Atg8*^78,79^ had no effect on protein intake (Figures S4A and S4B), indicating that the role of Lsp2 in protein homeostasis may be autophagy independent.

We next hypothesised that amino acid deprivation might be accompanied by Lsp2 proteolysis to liberate amino acids. Therefore, interfering with proteasome function in peptidergic neurons might maintain high Lsp2 levels and reduce perceived protein deprivation and yeast intake. Expressing RNAi against the *proteasome α7 subunit* (*Prosα7*), as well as thermo-sensitive dominant negative alleles of *Prosβ2* and *Prosβ6*^80^ under *dimm-GAL4* control had no effect on deprivation-induced yeast consumption (Figures S4C and S4D). We conclude that Lsp2-dependent regulation of protein intake may not be directly mediated by its degradation, and instead could rely on direct sensing of Lsp2 levels by these neurons.

Larval serum proteins are synthesised in large quantities by the fat body and circulate in the haemolymph^69,81,82^. To test whether cellular uptake of circulating Lsp2 could regulate protein appetite, we knocked down *Fbp1*^81^, which encodes a candidate receptor mediating larval serum protein endocytosis, in peptidergic neurons. This produced no detectable effect on protein consumption, suggesting that protein appetite is controlled by variations in the intracellular Lsp2 pool, rather than the ability of peptidergic neurons to import circulating Lsp2.

### Lsp2 regulates protein feeding through specific peptidergic populations

To determine if a specific subtype of peptidergic cells is responsible for the effect of *Lsp2* on protein homeostasis, we knocked down *Lsp2* in individual peptidergic subtypes, using a range of targeted Gal4 drivers. Subtypes were chosen for their known roles in feeding regulation, and/or stronger *Lsp2* downregulation in our dataset. In most tested cells, this intervention did not alter yeast consumption (Figures S5A-S5E). Surprisingly, targeting *Lsp2* in neurons expressing *Diuretic hormone 31* (*Dh31*) reduced yeast consumption (Figures 5A and 5B), in contrast to the increase observed with pan-peptidergic knockdown. This suggests that the output of *Dh31*+ neurons might be tied to intracellular amino acid availability rather than *Lsp2* levels: increased free amino acids in *Lsp2* knock-downs could promote Dh31-related signals to enhance feeding suppression. The same manipulation in neurons expressing *Myoinhibiting peptide precursor* (*Mip*, also known as *Allatostatin B*) boosted yeast consumption after protein deprivation (Figure 5C), phenocopying pan-peptidergic knock down. Unexpectedly, these flies also displayed increased yeast appetite when sated (Figures 5E). In both cases these effects were absent from genetically identical temperature controls maintained at the restrictive temperature (Figures 5D and 5F). These results suggest that *Lsp2* downregulation modulates protein appetite through distinct effects across peptidergic subtypes.

**Figure 5.**
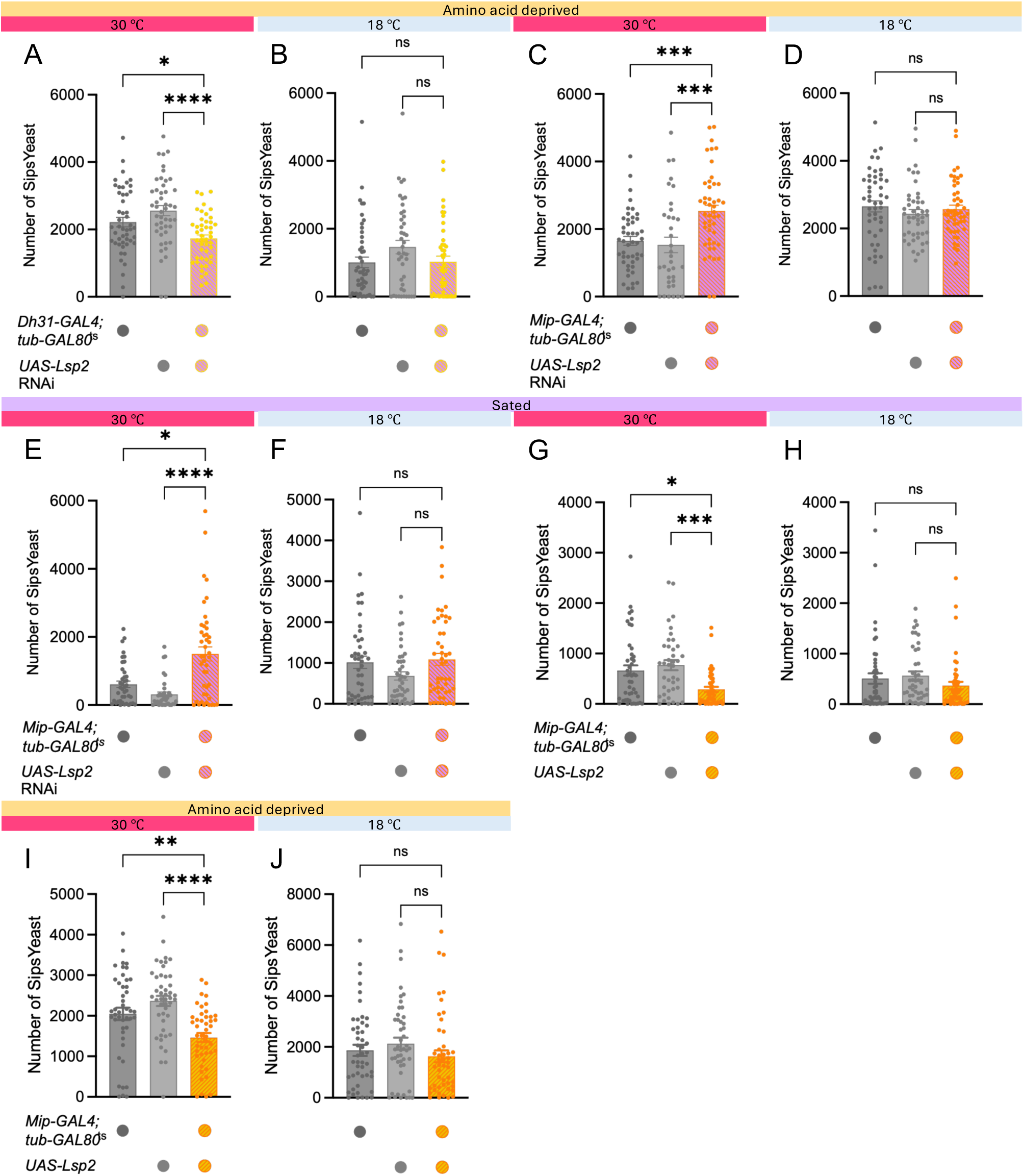
Lsp2 levels in *Mip*+ and *Dh31*+ neurons regulates yeast intake (**A-B**) Reduced yeast intake was measured in amino acid deprived flies with *Lsp2* RNAi expressed in *Dh31*+ neurons (A) but not in genetically identical controls kept at 18°C (B). n = 42-48 flies per group. (**C-D**) Increased yeast intake was measured in amino acid deprived flies with *Lsp2* RNAi expressed in *Mip*+ neurons (C) but not in genetically identical controls kept at 18°C (D). n = 35-48 flies per group. (**E-F**) Increased yeast intake was measured in sated flies with *Lsp2* RNAi expressed in *Mip*+ neurons (E) but not in genetically identical controls kept at 18°C (F). n = 37-47 flies per group. (**G-H**) Reduced yeast intake was measured in sated flies with *Lsp2* overexpressed in *Mip*+ neurons (G) but not in genetically identical controls kept at 18°C (H). n = 39-48 flies per group. (**I-J**) Reduced yeast intake was measured in protein deprived flies with *Lsp2* overexpressed in *Mip*+ neurons (I) but not in genetically identical controls kept at 18°C (J). n = 45-48 flies per group. ns p ≥ 0.05, * p < 0.05, ** p < 0.01, *** p < 0.001, **** p < 0.0001, Kruskal-Wallis ANOVA with Dunn’s multiple comparisons test. Individual data points are single flies. Barplots are mean +/- standard error of the mean (SEM).

We hypothesised that increased Lsp2 levels within *Mip*+ cells may mimic a state of protein satiety. To test this, we generated a *UAS-Lsp2* strain allowing cell-specific overexpression. Consistent with a bidirectional role for Lsp2 in protein appetite regulation, overexpressing *Lsp2* in *Mip*+ neurons reduced yeast consumption in both amino acid deprived and sated flies (Figures 5G and 5I), but not in genetically identical flies maintained at the restrictive temperature (Figures 5H and 5J). These results suggest that Lsp2 levels within *Mip*+ neurons act in a homeostatic manner to encode protein demand, and highlight *Mip*+ neurons as a key regulatory node in protein feeding.

### Monitoring Lsp2 dynamics under protein hunger

To assess whether transcriptional regulation of *Lsp2* is reflected at the protein level and investigate its spatial organisation within *Mip*+ neurons, we generated flies carrying an endogenous GFP tag on the C-terminal end of Lsp2. Imaging intact heads under a fluorescence microscope revealed broad GFP signal in the proboscis, eyes, and other head structures, consistent with Lsp2 being abundant in the circulating hemolymph and widely distributed across tissues (Figures S5H and S5I), although fluorescence measured in the proboscis remained unchanged under amino acid deprivation (Figure S5J). In the brain, simultaneous labelling of *Mip*+ neurons enabled direct visualisation of Lsp2 within these cells (Figure 6A), revealing that the protein is organised into discrete puncta scattered throughout the cytoplasm (Figure 6B). Consistent with the reduction in transcript levels, Lsp2 protein abundance in *Mip*+ neurons decreased following three days of amino acid deprivation (Figure 6C), reflected both in a general reduction in fluorescence intensity (Figure 6D) and a decreased number of puncta (Figure 6E).

**Figure 6.**
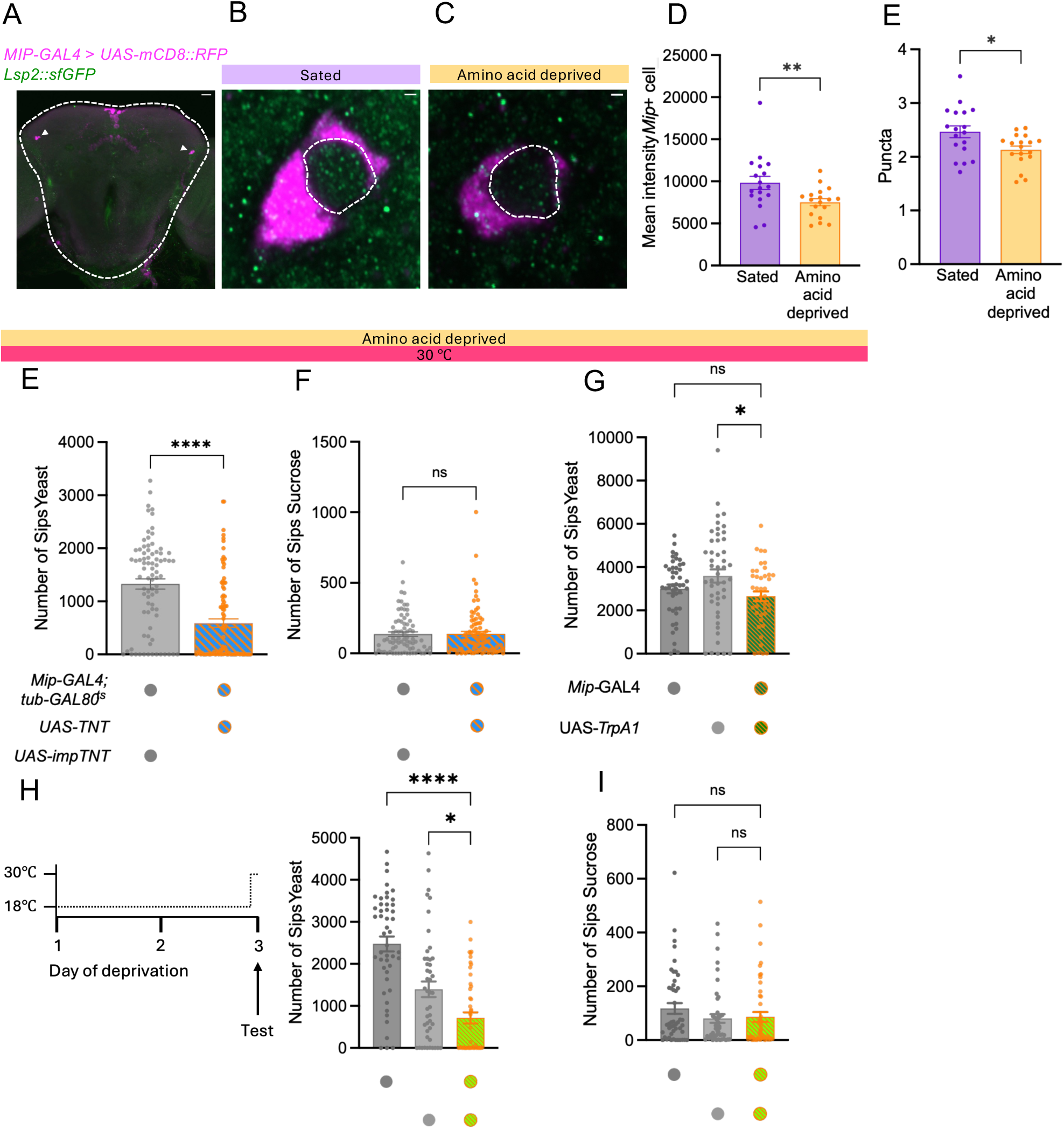
*Mip*+ neurons control protein intake (**A**) RFP (magenta) labelling of *Mip*+ neurons with Lsp2 protein labelled with GFP (green) in a *Drosophila* central brain. Scale bar is 1µm. (**B-C**) Representative *Mip*+ somas (magenta) with Lsp2 puncta (green), in sated (B) and amino acid deprived (C) fly brains. The area used for fluorescence quantification is outlined. Scale bar is 10µm. (**D-E**) Quantification of mean fluorescence intensity (D) and puncta numbers (E) in *Mip*+ neurons from sated and amino acid deprived brains. n = 18 flies per group. (**E-F**) Reduced yeast intake (E) but not sugar intake (F) was measured in amino acid deprived flies expressing *TNT* in *Mip*+ neurons, in comparison to controls expressing impaired *TNT*. n = 78-91 flies per group. (**G**) No significant change in yeast consumption was measured in amino acid deprived flies when *Mip*+ neurons were continuously activated with Trpa1 during the deprivation and test phases. n = 59-64 flies per group. (**H-I**) Reduced yeast intake (H) but not sucrose intake (I) was measured in amino acid deprived flies when *Mip*+ neurons were activated with Trpa1 during the test phase only. n = 46-48 flies per group. ns p ≥ 0.05, * p < 0.05, ** p < 0.01, **** p < 0.0001, Kruskal-Wallis ANOVA with Dunn’s multiple comparisons test. Individual data points are single flies. Barplots are mean +/- standard error of the mean (SEM).

### *Mip*+ neurons control protein intake

Single-cell RNA-seq and RT-qPCR experiments (Figure S5F) did not detect any changes in *Mip* expression under protein deprivation, suggesting that protein hunger does not involve altered Mip transcription. We also demonstrated that *Mip* transcripts are not directly influenced by Lsp2 knock down (Figure S5G). Production of potentially co-transmitted fast-acting neurotransmitters^45,83,84^ is also likely unaffected, as none of the genes regulating their production were differently expressed under protein deprivation. Instead, Lsp2 fluctuations might modulate signal release from *Mip*+ neurons. To address the role of *Mip*+ neurons in protein hunger, we blocked synaptic exocytosis by expressing *tetanus toxin light chain* (*TNT*)^85^. This inhibition during the deprivation and test periods decreased yeast consumption, in comparison to controls expressing the inactivated *TNT* gene (*impTNT*; Figure 6E) without altering sucrose intake (Figure 6F). This phenotype was not visible in genetically-identical flies kept at permissive temperature (Figure S6A).

To address whether activation of *Mip*+ cells also influenced yeast consumption, we ectopically expressed *TrpA1*, a temperature-sensitive Ca^2+^ channel that depolarises neurons at temperatures above 25°C^86^. Chronic activation of *Mip*+ cells during the deprivation and test phases did not alter feeding behavior (Figure 6G); however, acute activation of *Mip*+ neurons during the feeding test only resulted in a failure of protein homeostasis (Figure 6H), without affecting sucrose intake (Figure 6I). This effect was absent from genetically identical temperature controls maintained at 18°C (Figure S6B). Together, these data suggest that bidirectional perturbation of *Mip*+ neuronal activity disrupts protein homeostasis, indicating that tightly regulated signalling from these neurons is required for the adaptive control of protein intake.

### Mip signalling functions independently of known protein hunger circuits

We next explored whether signals from *Mip*+ neurons interact with known circuits controlling protein hunger. Notably, protein deprivation increases the activity of a pair of PPM2 dopaminergic neurons to trigger yeast preference via activation of a single neuron that projects to the fan-shaped body and lateral accessory lobe (FB-LAL)^17^. Mip peptides are ligands of the sex peptide receptor (SPR)^87,88^, and *in vitro* evidence also suggests they can bind the CCHamide-1 receptor (CCHa1-R)^89^. We knocked down each of these receptors in both FB-LAL (Figures S6C-S6F) and dopaminergic (Figures S6G-S6J) neurons during protein deprivation, but observed no effect on yeast consumption. *Mip* is expressed in a sparse population of ∼50-70 neurons, scattered through the CNS, and innervating areas of the brain including the antennal lobe and suboesophageal zone, two structures essential for chemical sensing of food cues^90,91^. We therefore wondered whether *Mip*+ neurons might instead influence protein appetite by modulating sensory detection of yeast cues. We knocked down both receptors in *Ir76b*-expressing neurons, which play essential roles in gustatory detection of amino acids and yeast-derived signals^2,15,20^. However, this manipulation also failed to alter yeast consumption (Figure S6K-L), suggesting that Mip-dependent regulation of protein appetite is unlikely to act through these known sensory pathways.

## Discussion

### Peptidergic neurons control protein homeostasis

Nutrient balance is a major determinant of animal behaviour. In *Drosophila*, the macronutrient composition of food influences behaviours as diverse as mating^92^, sleep structure^93^ and feeding^15,17^. Physiologically, depletion of specific nutrients seems embodied in gene expression signatures dissimilar from that of general starvation^73^. Our findings demonstrate that some peptidergic neurons are tuned to regulate protein homeostasis behaviour, and provide a readout of the gene expression landscape in these cells following amino acid deprivation.

Peptidergic signalling conveys protein satiety signals for the cessation of feeding^29,94^. To our knowledge, our work represents the first demonstration that peptidergic signalling also controls compensatory yeast intake following amino acid deprivation. Of the diverse peptidergic cell classes, *Mip*+ cells may be particularly important for protein homeostasis. In contrast to previous findings describing Mip’s role in general hunger^91^, both chronic inhibition and transient activation of these neurons impair protein, but not sugar homeostasis behaviour, suggesting that balanced activity of *Mip*+ neurons is required for scaling protein consumption in line with internal nutrient requirements.

Recording calcium transients and changes in Mip release over the protein deprivation period will help determine the precise mechanism(s) by which *Mip*+ cells encode protein deficiency. The fact that these neurons may also comprise a functionally heterogeneous population is important to keep in mind^90,95^. For example, decreased female receptivity post-mating might be controlled by dorsal *Mip*+ neurons only^96^. Further efforts to delineate specific roles for individual *Mip*+ subpopulations will be valuable. Relatedly, our study focused specifically on mated females, which leaves open questions about the role of Lsp2 and Mip in male flies, and how mating status might influence Lsp2 control of protein appetite.

### Lsp2 abundance acts as a gauge to determine appropriate protein consumption

Under amino acid deprivation, peptidergic neurons in the fly brain show widespread downregulation of gene expression, likely with the objective of preserving amino acid resources. However, *Lsp2* downregulation also plays a role in determining the extent of yeast refeeding following deprivation. The consequences of *Lsp2* modulation are reciprocal, with *Lsp2* knockdown in *Mip*+ neurons resulting in protein overconsumption, while overexpression results in underconsumption. Thus, some mechanism may exist in *Mip*+ neurons to determine appropriate homeostatic responses depending on a read out of Lsp2 levels.

The mechanism of action of Lsp2 is not clear, however. We show that preventing proteolysis or autophagy did not alter protein intake, which indicates that the availability or cellular usage of Lsp2 degradation products are not required for downstream signalling from these cells to control protein feeding - although we cannot completely exclude that our manipulations were insufficient to adequately suppress Lsp2 breakdown. One alternative hypothesis is that Lsp2 could function as part of a signalling cascade encoding protein hunger. Initially believed to only function as an amino acid storage during development, mounting evidence now proposes broader roles for Lsp proteins across adult tissues^69^. In particular, Lsp2 promotes translation initiation by acting on 4E-BP phosphorylation and amino acid incorporation into ribosomal proteins, a process mediated by the amino acid content of larval diet^70,71^. It is therefore plausible that Lsp2 also interacts with translational pathways in *Mip*+ cells, potentially influencing the production of Mip peptides or regulators of peptide or neurotransmitter release.

Physiological, transcriptional and behavioural effects of protein deprivation depend on the specific amino acids that are absent from the diet, reflecting the distinct roles of individual amino acids as signalling molecules and neurotransmitter precursors^97–99^. Lsp2 is enriched in aromatic amino acids, including tyrosine, whose restriction alone reproduces Lsp2-associated effects on translational suppression and longevity^71,100^. Therefore, Lsp2 function may be linked to specific amino acid availability rather than general protein depletion. Defining which amino acid deprivation states are required for Lsp2-dependent control of protein homeostasis will also help resolve underlying mechanisms.

## Data and resource availability

Single-nucleus RNA-seq data is publicly available on GEO (GSE334747). Code for snRNA-seq analysis is available at https://github.com/vincr04/Lsp2_2006/. Requests for further information, code, resources, or reagents should be directed to Vincent Croset.

## Acknowledgements

We thank the Vienna *Drosophila* Resource Center and Bloomington *Drosophila* Stock Center for flies. We thank Helen Siddle, Jaryd Mercer, Molly Amlot and Lucas Yan for laboratory assistance, Nicola Fullard and Cole Sims for help with FACS, and Rafiqul Hussain and Johnathan Coxhead from Newcastle GCF for their support with single-nucleus RNA-seq. We are grateful to James Evans, Alasdair Boeddinghaus, and members of the Croset group for discussion, and Chris Allen, Wolf Huetteroth, and Gaurav Das for their comments on the manuscript. This work was supported by the UK Biotechnology and Biological Sciences Research Council (BB/W007347/1) and starting funds from Durham University.

## Author contributions

Designed research S.W., V.C.; Performed research S.W., Y.L., V.C.; Analysed data S.W., Y.L., V.C.; Resources S.W., V.C.; Writing S.W., V.C; Supervision V.C.; Funding acquisition V.C.

## Declaration of interests

The authors declare no competing interests

## Declaration of generative AI in the writing process

During the preparation of this work, the authors used GPT-5.2 as a language support tool to rephrase certain sentences and clarify the wording of specific statements. A pre-submission review was conducted using q.e.d. Science. All scientific content, interpretation, and final text were produced and verified by the authors, who take full responsibility for the content of the publication.

## STAR * Methods

### KEY RESOURCE TABLE

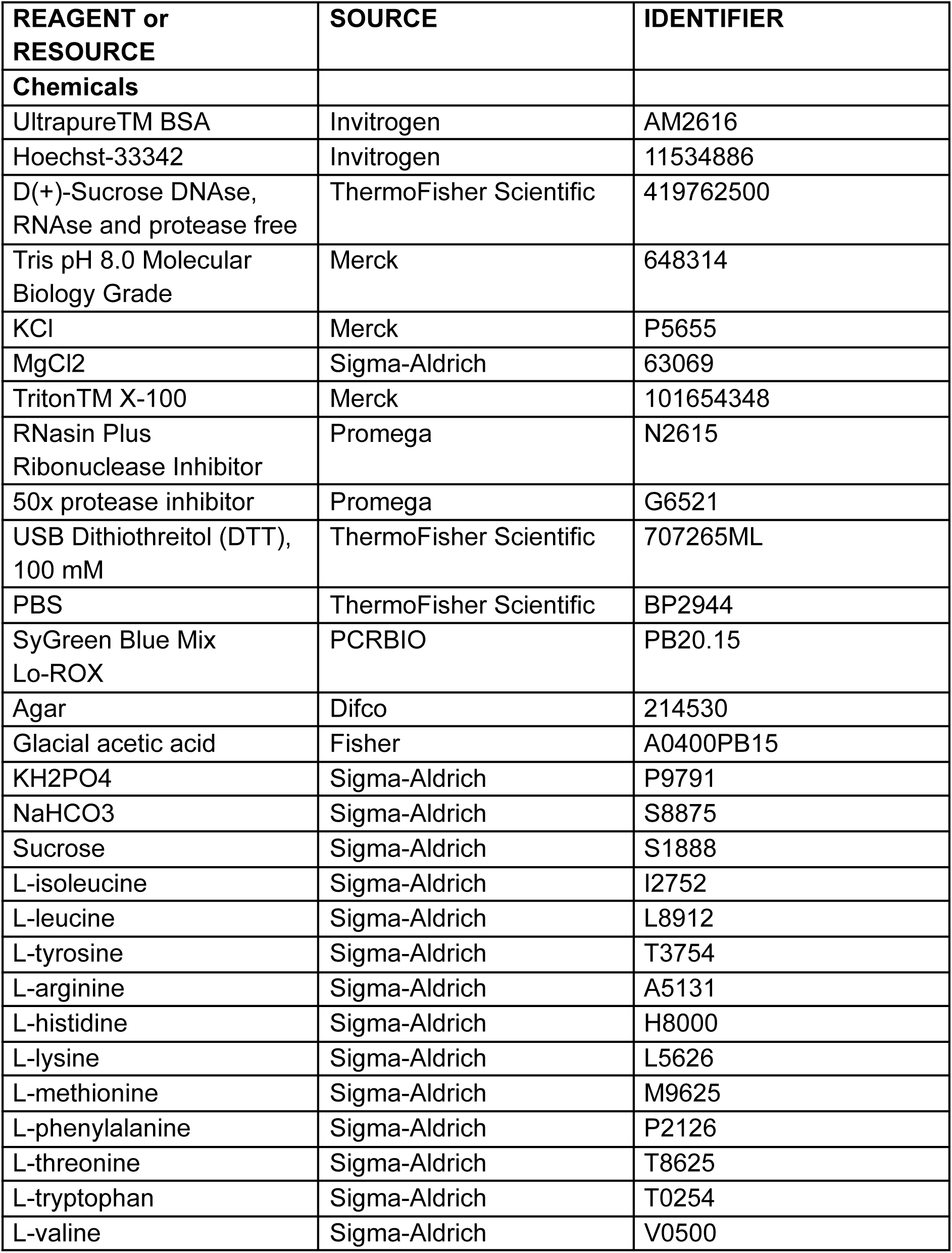
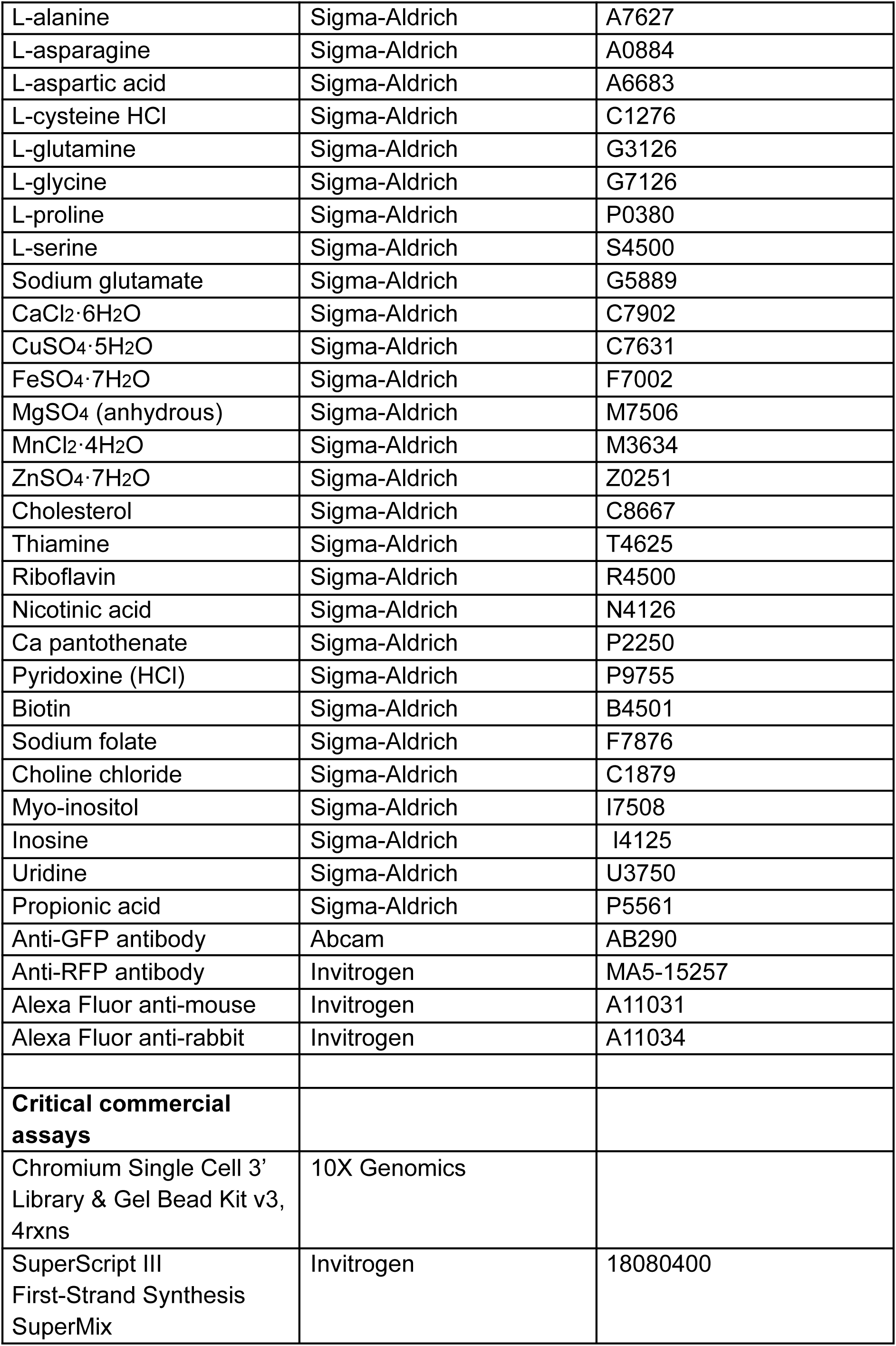
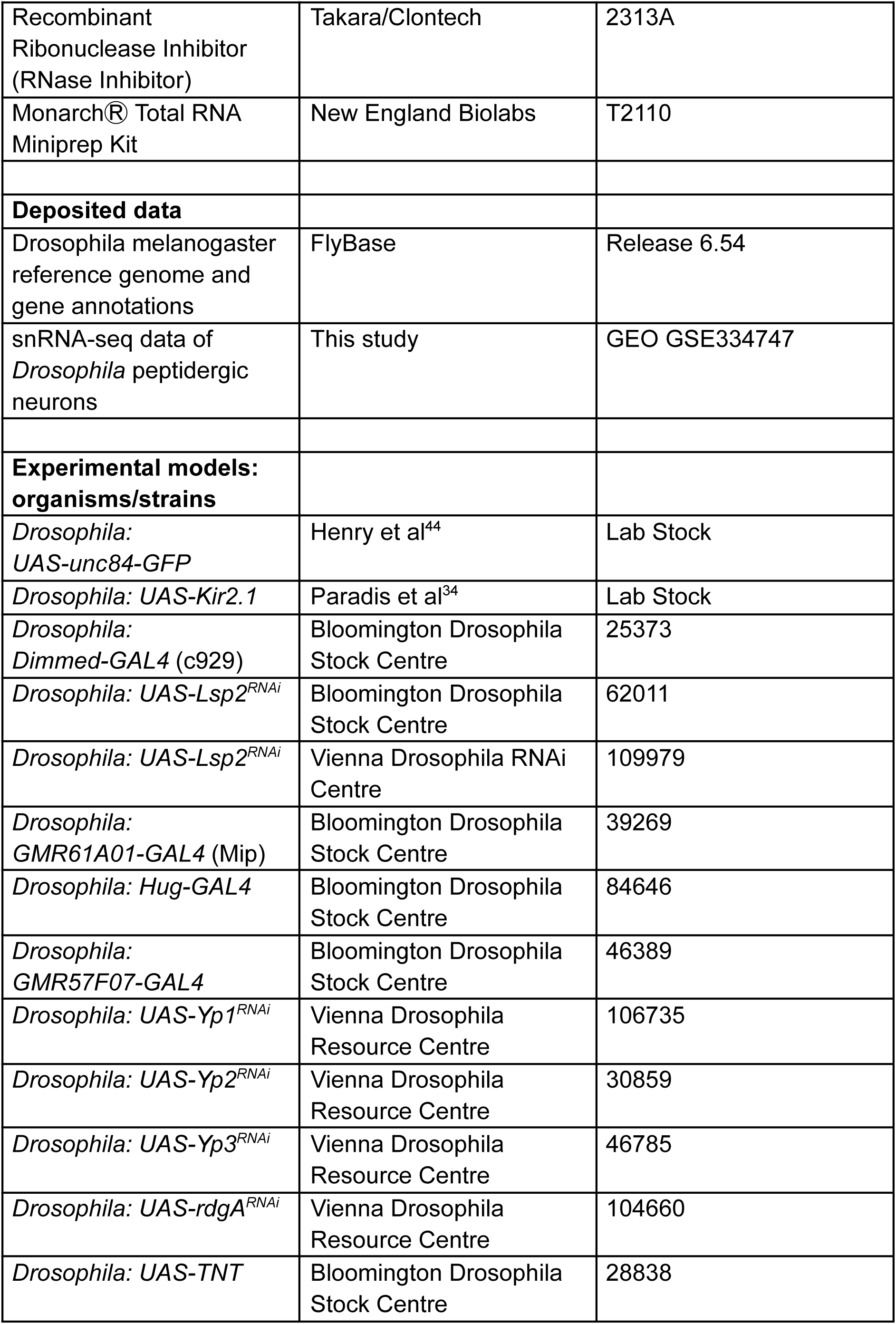
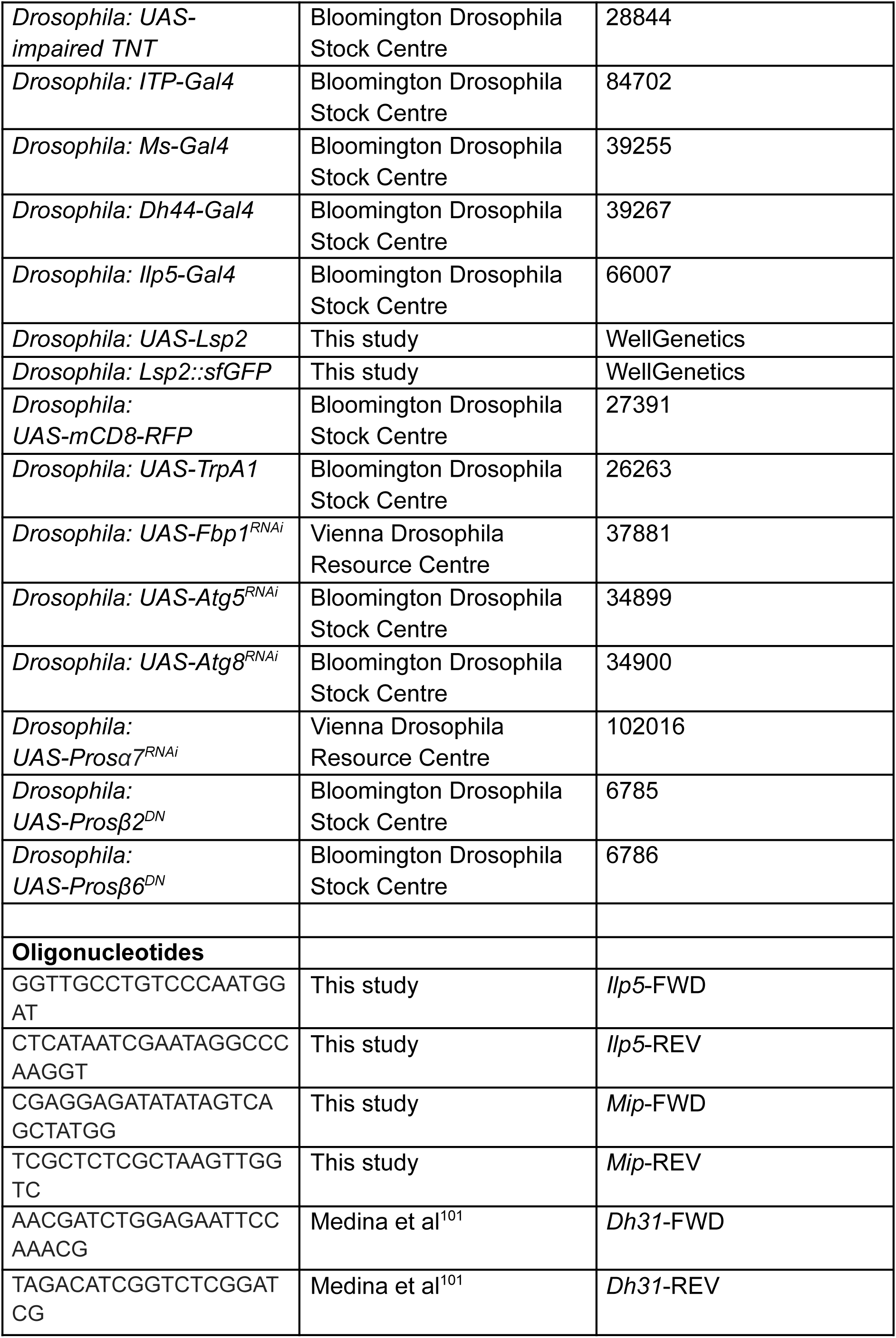
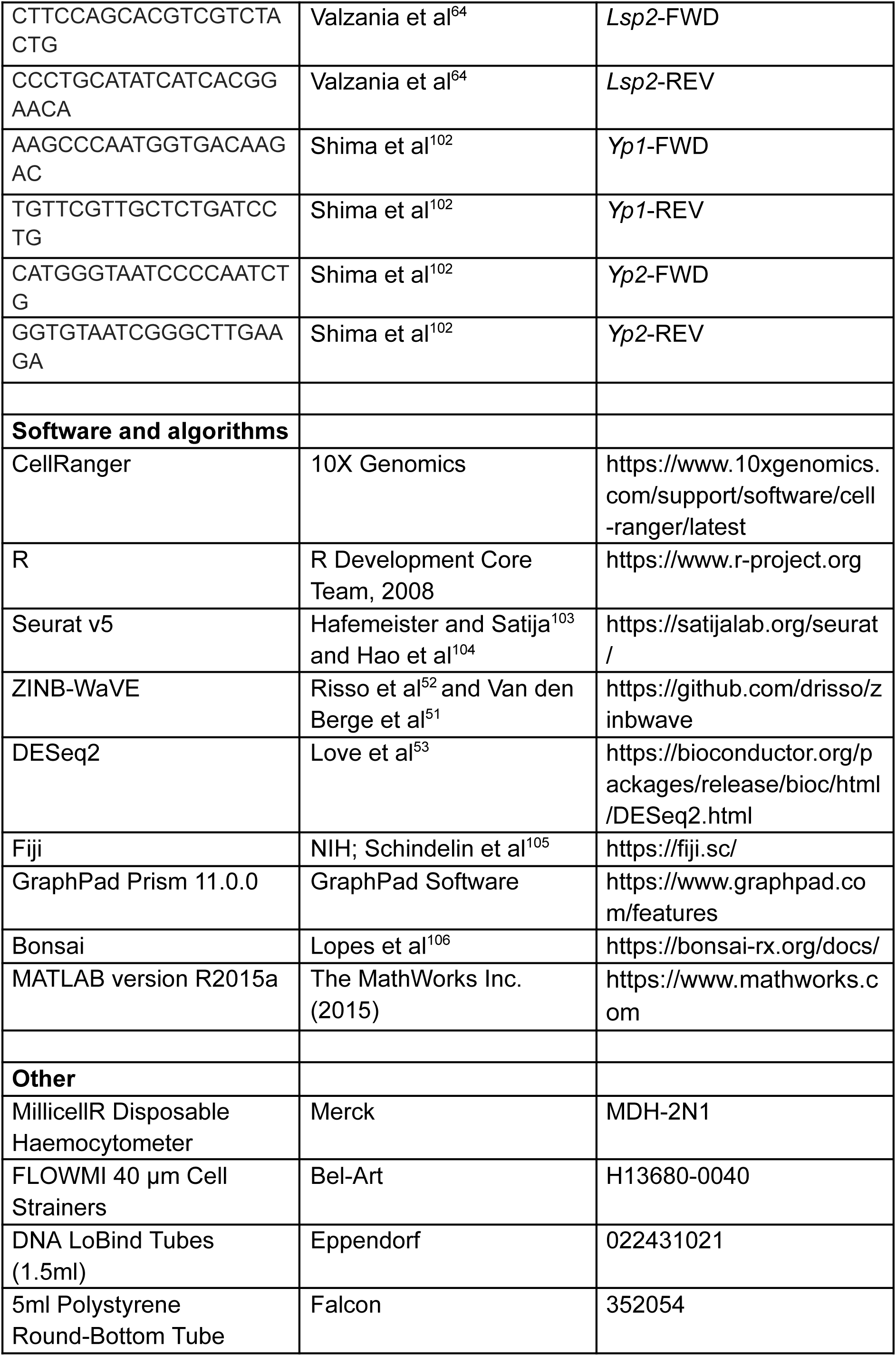
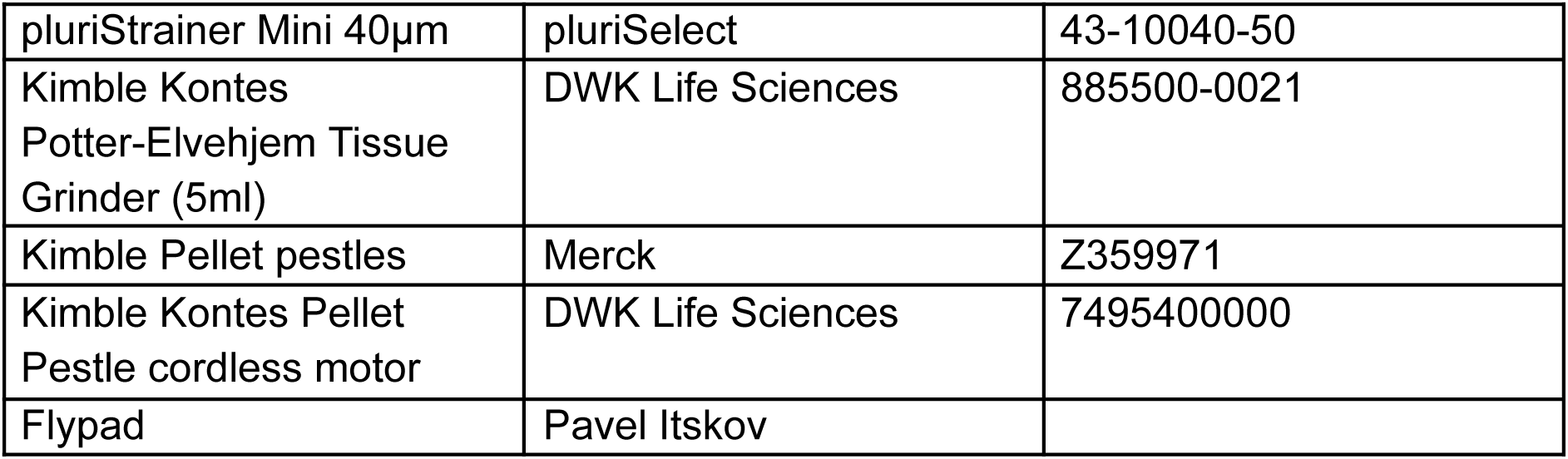

### Experimental model and subject details

#### Drosophila husbandry

Flies were raised on standard cornmeal food under a 12:12 light:dark cycle at 60% humidity and 25°C unless otherwise stated. 6-10 day old female flies were used for nutrient homeostasis experiments, RT-qPCR, imaging and single-nuclei transcriptomics. Chemically-defined (holidic) diet was prepared as outlined in Piper 2014^11^, using exome-matched amino acid ratios^11^. Gal4 drivers used in this study are dimm-GAL4 (BDSC 25373), GMR61A01-GAL4 (BDSC 39269), Hug-GAL4 (BDSC 84646), ITP-GAL4 (BDSC 84702), Ms-GAL4 (BDSC 39255), Dh44-GAL4 (BDSC 39267), Ilp5-GAL4 (BDSC 66007). UAS lines are UAS-unc84-GFP^44^, UAS-Kir2.1^34^, UAS-Lsp2 RNAi (BDSC 62011, VDRC 109979), UAS-Yp1 RNAi (VDRC 106735), UAS-Yp2 RNAi (VDRC 30859), UAS-Yp3 RNAi (VDRC 46785), UAS-rdgA RNAi (VDRC 104660), UAS-TNT (BDSC 28838), UAS- impaired TNT (BDSC 28844), UAS-*Lsp2* (this study), UAS-mCD8-RFP (BDSC 27391), UAS-TrpA1 (BDSC 26263), UAS-Fbp1RNAi (VDRC 37881), UAS-Atg5 RNAi (BDSC 34899), UAS-Atg8 RNAi (BDSC 34900), UAS-Prosα7 (VDRC 102016), UAS-Pros□2 (BDSC 6785), UAS-Pros□6 (BDSC 6786).

#### Generation of transgenic *Drosophila*

Transgenic flies carrying UAS-Lsp2 were generated by WellGenetics Inc. (New Taipei City, TW) as follows: a 2152 bp genomic fragment containing the Lsp2 coding sequence was amplified by PCR from w[1118] genomic DNA, with a 4 bp Drosophila Kozak sequence added immediately upstream of the start codon. The amplified fragment was cloned into the XhoI site of the pUAST-attB vector using sequence-and ligation-independent cloning (SLIC)^107^. The resulting pUAST-attB-Lsp2 construct was integrated at the VK00027 attP landing site on chromosome 3R (89E11)^108^ by embryo microinjection.

Insertion of a sfGFP-3xP3-RFP cassette at the C-terminus of Lsp2 was performed by WellGenetics Inc. by CRISPR-mediated mutagenesis, using a modified method from ^109^. In brief, DNA plasmids containing the *Lsp2*/*CG6806*-targeting gRNA sequence GTCCAGGATCTAGACCACAT[TGG], donor template with sfGFP, a floxed 3xP3-RFP, and two homology arms, and *hs-Cas9* were microinjected into w[1118] embryos. Insertions in F1 flies carrying the 3xP3-RFP selection marker were further validated by genomic PCR and sequencing.

### Method details

#### FlyPAD assays

Around 3-5 days post-eclosion mated females were sorted into vials of holidic *Drosophila* medium. After a 72 hour habituation period, flies were transferred to either a fresh holidic medium vial or holidic medium without amino acids, and were maintained on this diet for a further 72 hours. Yeast and sucrose consumption were assessed using FlyPAD, as described^30^. One FlyPAD food well was filled with 5% sucrose and the other 5% yeast, each in 1% agarose. Flies were individually transferred to FlyPAD arenas and allowed to interact with food sources for one hour. FlyPAD data were acquired using custom-written Bonsai software^106^ and analysed using custom-written MATLAB programmes as described^30^. Flies were allocated and transferred to FlyPAD consoles in an order counterbalanced for dietary condition and genotype. For thermogenetic experiments flies were raised at 18°C and transferred to 30°C for the three days deprivation period and FlyPAD measurement, unless otherwise specified. Behaviour data were analysed using GraphPad Prism.

Depending on normality, one-way ANOVA with Dunnett’s multiple comparisons or Kruskal-Wallis ANOVA with Dunn’s multiple comparisons were used. Significance was defined using alpha < 0.05.

#### Single nuclei sample preparationh

Groups of 2000 mated female flies underwent feeding and deprivation as described above, split in vials of 50. After 72 hours in their dietary conditions, flies were flash-frozen in liquid nitrogen to extract whole heads, which were processed immediately or stored at -80°C.

Nuclei were prepared as described^110^. Briefly, samples were mechanically dissociated for 30s in 1ml homogenisation buffer (250 mM Sucrose, 10 mM Tris-HCl pH 8.0, 25 mM KCl, 5 mM MgCl2, 0.1% Triton-x 100, 0.5% RNasin Plus, 1X 50X protease inhibitor, 0.1 mM DTT) using a pestle motor (DWK Life Sciences, 7495400000) before being released by 25 strokes of a dounce pestle (DWK Life Sciences, 8855000021). These steps were performed on ice. The homogenate was filtered through a 40µm cell strainer (pluriSelect, 43-10040-50) to remove unlysed tissue and again using a 40µm Flowmi (BelArt, H13680-0040) into a 1.5ml Eppendorf tube. Samples were centrifuged at 1000 x g for 10 mins at 4°C, supernatant discarded and pellet resuspended in 500µl resuspension buffer (1X PBS, 0.5% BSA, 0.5% RNasin Plus). Samples were filtered once more using 40µm Flowmi into a 5ml FACs tube and incubated with Hoechst-33342 (1:1000, Invitrogen, 11534886) for 15 minutes on ice prior to fluorescent sorting.

#### Fluorescent nuclei sorting

Fluorescent sorting of nuclei was performed on a BD FACS Melody (BD Biosciences) using the 405nm, 488nm and 640nm lasers and a 100µm nozzle. Gating was achieved by selecting Hoechst-33342- and GFP-positive nuclei. A final gate using FSC-H and FSC-A was used to prevent collection of clumps or doublets (Figure S2B). Each sample was sorted into a collection tube with 500µl of resuspension buffer maintained at 4°C, and kept on ice prior to 10X barcoding. For each 10X Genomics run ∼20,000-50,000 nuclei were collected. Nuclei were spun for 10 minutes at 1000 x g at 4 and resuspended in 20 µl resuspension buffer to obtain concentrations of ∼800-1000 nuclei/µl. 1µl of the final suspension was diluted 10fold and used for a haemocytometer to estimate sample concentration.

#### Library preparation, sequencing, preprocessing and quality control

mRNA barcoding was performed using 10X Genomics Chromium Single Cell 3’ Reagent Kit v3 according to manufacturer’s instructions. Two libraries were prepared from nuclei in the sated condition, and one from nuclei in the amino acid deprived condition. Libraries were sequenced with NovaSeq 6000 (Illumina) at Newcastle Genomics Core Facility. We obtained on average 48,477 reads per cell and aligned these to the FB2023_06 *Drosophila melanogaster* genome assembly (v6.52) using CellRanger 7.2.0 to create digital gene expression (DGE) matrices which were analysed further in Seurat v5^104^. Cell barcodes with <200 or >4000 features, >5% mitochondrial RNA, >20% ribosomal proteins and >20000 UMIs were removed. DGE matrices were then merged by condition, normalised and scaled.

#### Clustering and annotation

A two-step pipeline was used for clustering, with the aim of annotating clusters primarily via neuropeptide expression while considering overall similarities in gene expression. The FBgg0000179 (3/11/2025) list of neuropeptide genes was used as the feature set to perform PCA, DGE matrices were then integrated using CCA anchor-based methods and cells were clustered by a shared nearest neighbor modularity optimization based clustering algorithm^111^ with a resolution of 1.

Following this, a phylogenetic tree of all clusters was constructed in PCA space. Clusters on neighbouring branches with less than 10 protein-coding genes differently expressed between them (Wilcoxon signed-rank test, adjusted *p*<0.05) were grouped into a single cluster^50^. Clusters were then annotated based on neuropeptide expression, and top markers for each cluster were calculated (Data S1). We controlled that each sample was represented in each cluster (Figure S2D)

#### Differential gene expression

To account for zero inflation triggered by dropout events, cell specific weights were calculated with ZINB-WaVE^51,52^. DESeq2^53^ was then used to calculate differential gene expression between sated and amino acid deprived conditions, for each cluster. Genes with log2FC>1 and adjusted p-value <0.05 were considered to be differentially expressed.

#### RT-qPCR

Total RNA was extracted from groups of 50 fly heads using Monarch Total RNA Miniprep Kit (NEB, T2110). mRNA was retrotranscribed using the SuperScript III First-Strand Synthesis SuperMix (Invitrogen, 18080400) following manufacturers instructions. qPCR was performed in a CFX Duet Real-Time PCR System (Bio-Rad, 12016265). Each 20µl reaction contained 20 ng cDNA, 0.4 mM of each primer, and 10 µl SyGreen Blue Mix Lo-ROX (PCRBIO, PB20.15). Cycles ran as: 95°C, 2′; 40x [95°C, 5″; 60°C, 30″]. Quantification was performed with the comparative 2-ΔΔCt method^112^, using GAPDH and SdhA as housekeeping genes.

#### Immunohistochemistry and image acquisition

Flies were anaesthetised on ice and dissected in ice-cold PBST (PBS + 0.5% Triton-X) before being transferred to 4% paraformaldehyde in PBST and incubated for 30 min at RT. Samples were washed three times in PBST and blocked in Normal Goat Serum 5% in PBST for 2h at RT. Samples were then incubated in primary antibody solution for 48 hours with rotation at 4°C. Samples were then subjected to two 2-minute washes with PBST and three 30 minute washes with PBST at RT with rotation before being incubated with secondary antibody overnight at 4°C with rotation. Following washes as described above, brains were mounted with the posterior face upwards in Vectashield (Vector Laboratories, H-1000-10). Primary antibodies were Anti-GFP (Abcam, AB290), Anti-RFP (Invitrogen, MA5-15257).

Secondary antibodies were Alexa Fluor 568 anti-mouse (Invitrogen, A11031) Alexa Fluor 488 anti-rabbit (Invitrogen, A11034). Images were captured on a Zeiss LSM 800 using 10X, 20X or 63X objectives. Images were acquired at 0.017 ✕ 0.017 ✕ 1µm resolution with 16-bit depth, scan speed 6 and frame-by-frame scanning.

Fluorescent images of whole fly heads were obtained on a Zeiss Apotome using a 10X objective.

#### Image analysis

In Fiji a ROI was drawn around the cytoplasm of the *Mip*+ cells from a single slice in the centre of the Z stack of each cell body using the red channel. This was saved and used to measure intensity in the green channel from the corresponding image, where puncta were also manually counted. Intensity and puncta counts were divided by the area measured to account for differing cell sizes. From each brain the left and right hemisphere dorsal-most *Mip*+ cells were analysed and intensity/puncta counts were averaged to provide one data point per dissected brain.

## Notes

### Competing Interest Statement

The authors have declared no competing interest.

