## Supplementary figures for "Intracellular amino acid scarcity sensing tunes protein hunger"

Intracellular amino acid scarcity

sensing tunes protein hunger

### Figure S1

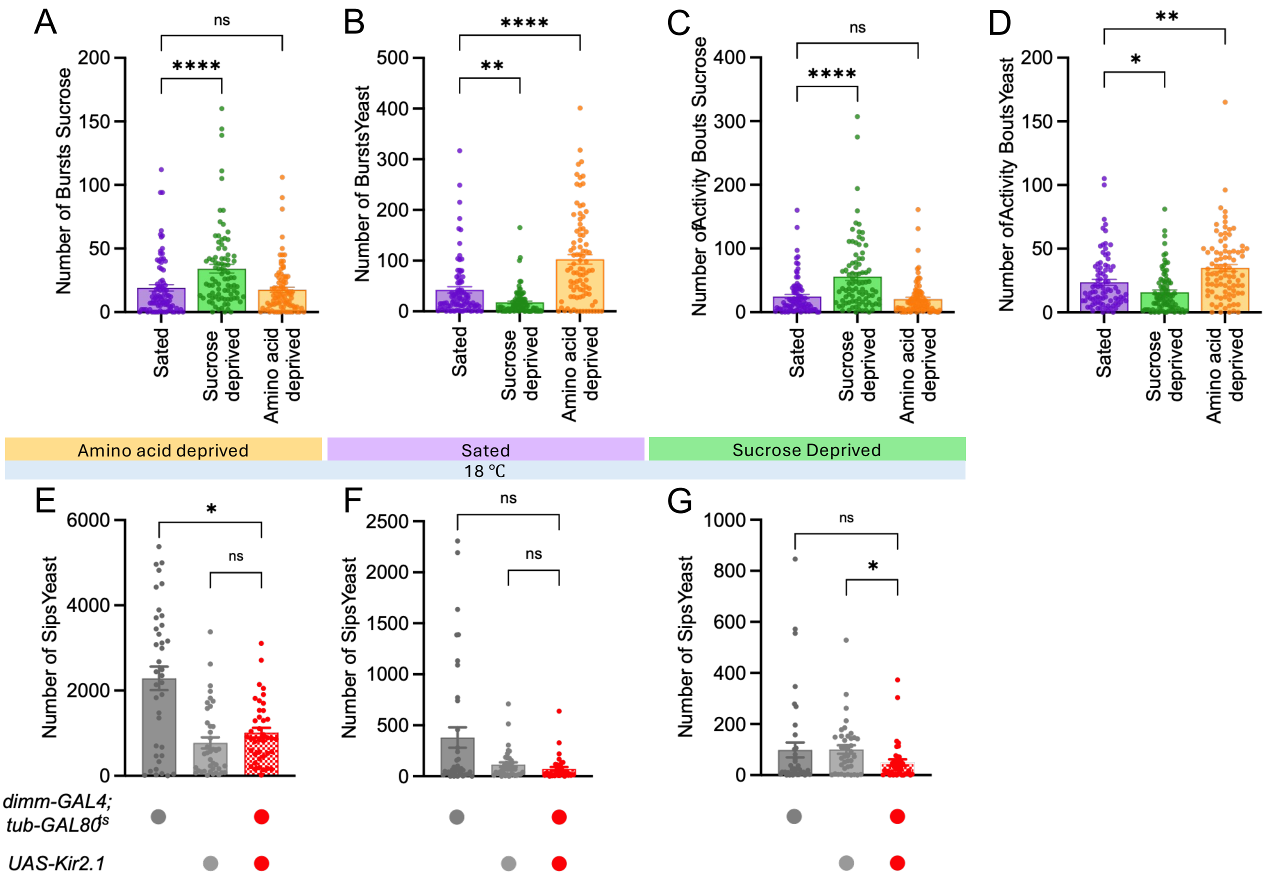

A
B
C
D
Number of Bursts Sucrose
Number of Bursts Yeast
Number of Activity Bouts Sucrose
Number of Activity Bouts Yeast
Sated
Sucrose deprived
Amino acid deprived
ns
****
**
*
Amino acid deprived
Sated
Sucrose Deprived
18 ℃
E
F
G
Number of Sips Yeast
dimm-GAL4; tub-GAL80ts
UAS-Kir2.1

Figure S1. Related to Figure 1.

(**A**) Number of feeding bursts on sucrose is increased in sucrose deprived flies in comparison to sated, but not amino acid deprived animals. n = 86-90 flies per group. (**B**) Number of feeding bursts on yeast is increased in amino acid deprived flies in comparison to sated and sugar deprived animals. n = 86-90 flies per group. (**C**) Number of activity bouts on sucrose is increased in sucrose deprived flies in comparison to sated, but not amino acid deprived animals. n = 86-90 flies per group. (**D**) Number of activity bouts on yeast is increased in amino acid deprived flies in comparison to and sugar deprived animals. n = 86-90 flies per group. (**E-G**) Flies carrying *dimm-Gal4*, *tub-GAL80$^{ts}$* and *UAS-Kir2.1* kept at permissive temperature (18ºC) consume similar amounts of yeast as genetic controls when amino acid deprived (E), sated (F), or sugar deprived (G). ns $p \geq 0.05$, * $p < 0.05$, ** $p < 0.01$, **** $p < 0.0001$, Kruskal-Wallis ANOVA with Dunn's multiple comparisons test. Individual data points are single flies. Barplots are mean +/- standard error of the mean (SEM).

### Figure S2

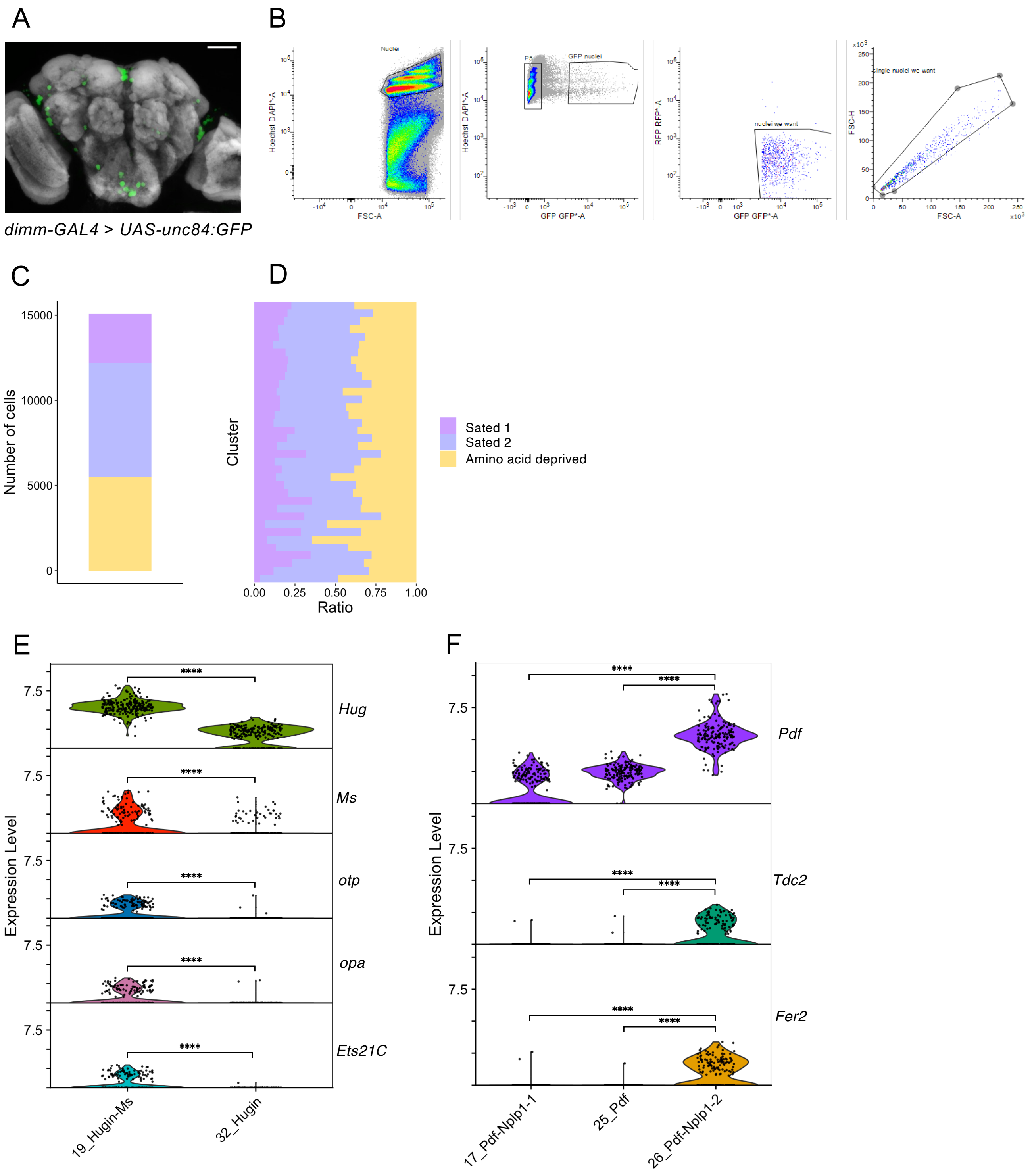
A
dimm-GAL4 > UAS-unc84:GFP
B
Nuclei
Hoechst DAPI*-A
FSC-A
P5
GFP nuclei
GFP GFP*-A
RFP RFP*-A
nuclei we want
single nuclei we want
FSC-H
C
Number of cells
15000
10000
5000
0
D
Cluster
Ratio
0.00
0.25
0.50
0.75
1.00
Sated 1
Sated 2
Amino acid deprived
E
Expression Level
7.5
****
Hug
Ms
otp
opa
Ets21C
19_Hugin-Ms
32_Hugin
F
Pdf
Tdc2
Fer2
17_Pdf-Nplp1-1
25_Pdf
26_Pdf-Nplp1-2

Figure S2. Related to Figure 2.

(**A**) unc84::GFP labelling of *dimmed-Gal4* nuclei in the Drosophila brain. Neuropil, labeled with nc82 anti-Brp antibody, is shown in grey. Scale bar is 20 µm. (**B**) FACS gating strategy to sort peptidergic nuclei. From left to right: Hoechst signal is used to distinguish nuclei from debris; GFP+ nuclei are sorted from GFP- nuclei; autofluorescent debris are removed from GFP+ nuclei with the RFP laser; singlets are identified by their uniform diameter. (**C**) Cumulative number of nuclei originating from each sample collected. (**D**) Ratio of nuclei originating from each sample, for each cluster. All clusters contain cells from each condition and sample. (**E-F**) Violin plots representing the expression of markers differently expressed between defined clusters of *Hugin*+ (E) and *Pdf*+ (F) neurons. **** $p < 0.0001$, Wald test with Benjamini-Hochberg adjusted p-values.

### Figure S3

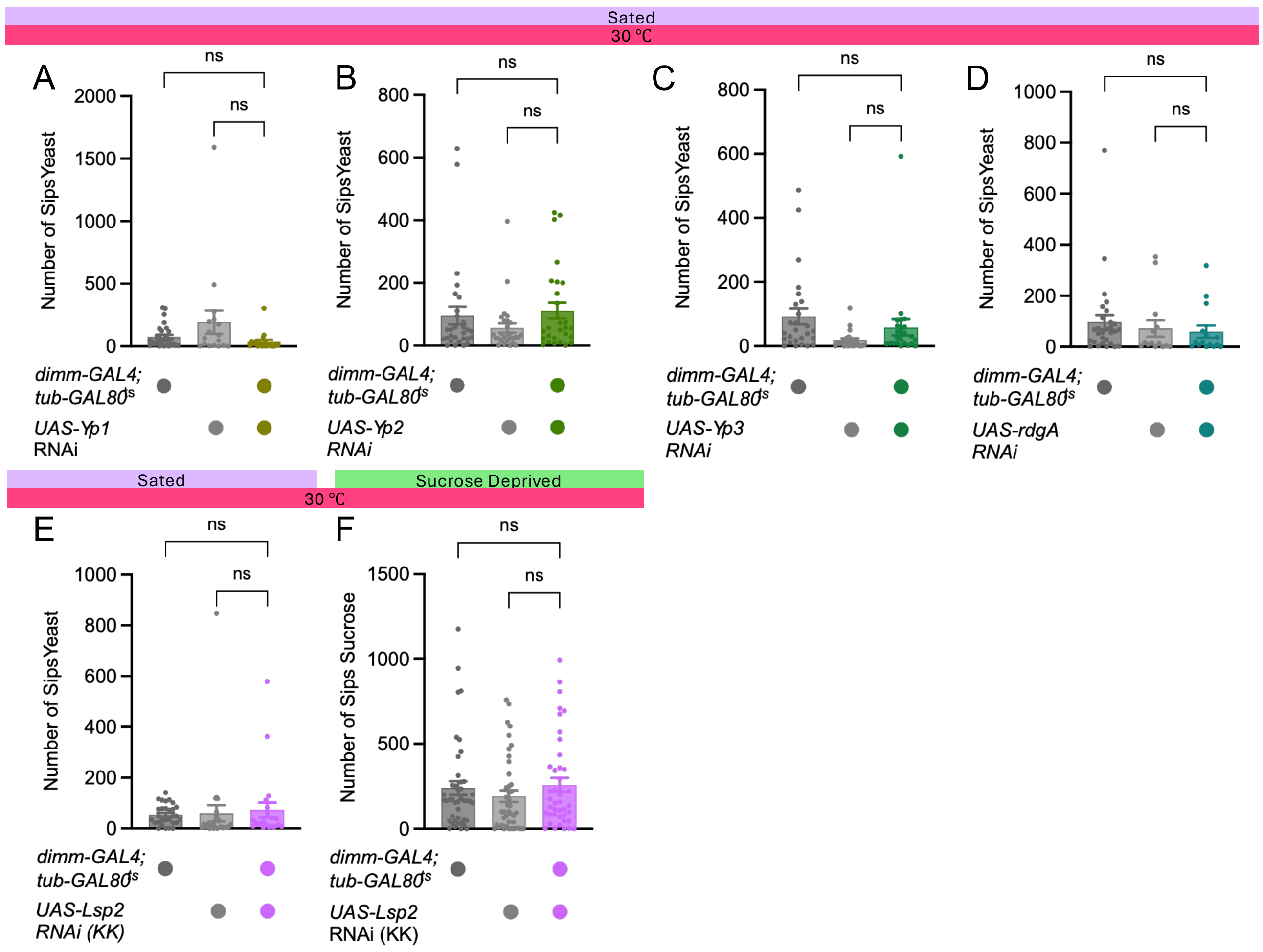

Sated
30 ℃
A
B
C
D
ns
ns
Number of Sips Yeast
dimm-GAL4; tub-GAL80ts
UAS-Yp1 RNAi
UAS-Yp2 RNAi
UAS-Yp3 RNAi
UAS-rdgA RNAi
Sated
Sucrose Deprived
30 ℃
E
F
Number of Sips Sucrose
UAS-Lsp2 RNAi (KK)

Figure S3. Related to Figure 4.

(**A-E**) No change in yeast intake was measured in sated flies expressing RNAi against *Yp1* (A), *Yp2* (B), *Yp3* (C), *rdgA* (D), and *Lsp2* (E) in peptidergic neurons using *dimmed-Gal4* driver. n = 14-29 flies per group. (**F**) No change in sugar intake was measured in sugar deprived flies expressing *Lsp2* RNAi in peptidergic neurons using *dimmed-Gal4* driver. n = 42-43 flies per group. ns $p \geq 0.05$, Kruskal-Wallis ANOVA with Dunn's multiple comparisons test. Individual data points are single flies. Barplots are mean +/- standard error of the mean (SEM).

### Figure S4

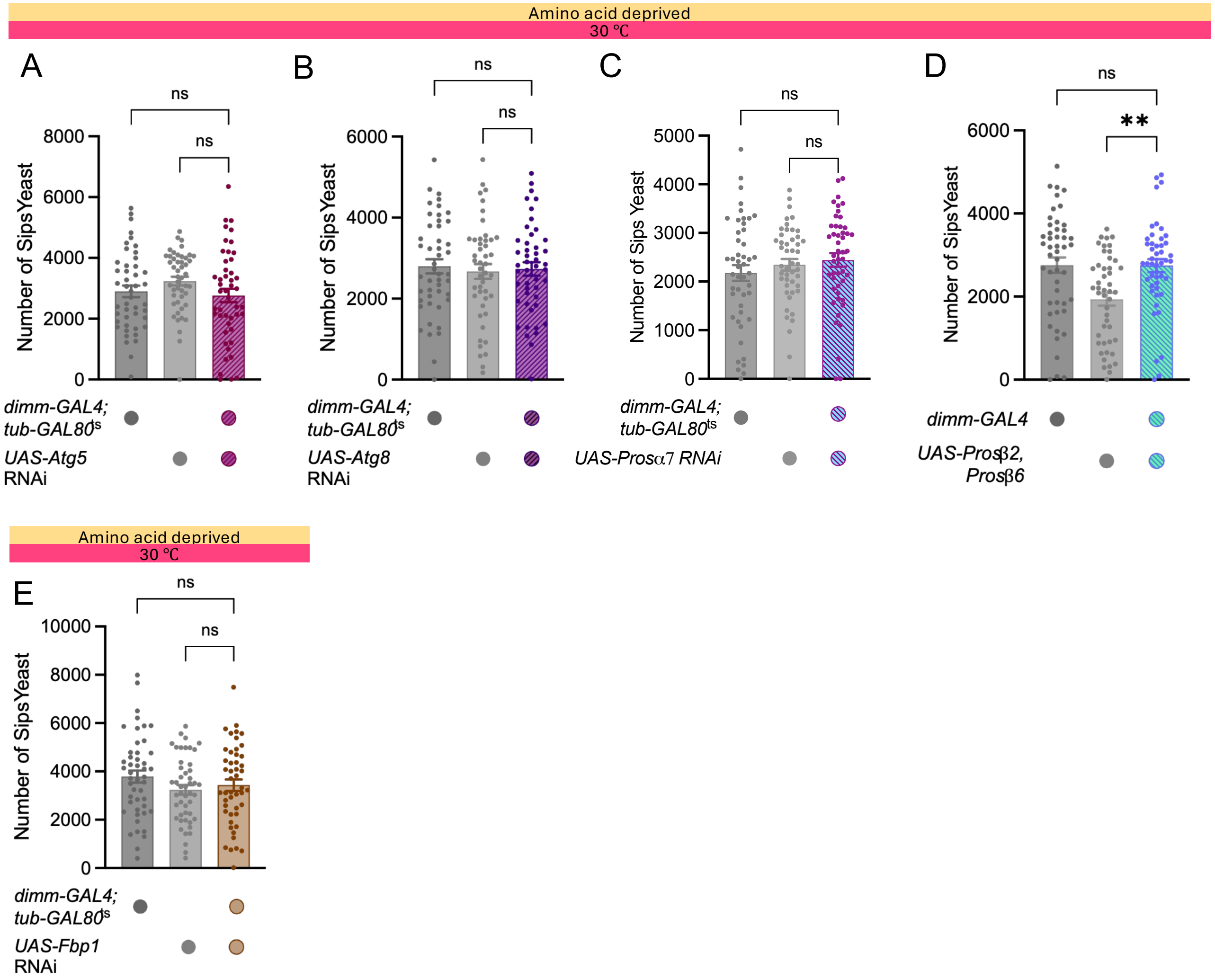
Amino acid deprived
30 ℃
A
ns
ns
8000
6000
4000
2000
0
Number of Sips Yeast
dimm-GAL4; tub-GAL80ts
UAS-Atg5 RNAi
B
ns
ns
6000
4000
2000
0
Number of Sips Yeast
dimm-GAL4; tub-GAL80ts
UAS-Atg8 RNAi
C
ns
ns
5000
4000
3000
2000
1000
0
Number of Sips Yeast
dimm-GAL4; tub-GAL80ts
UAS-Prosα7 RNAi
D
ns
**
6000
4000
2000
0
Number of Sips Yeast
dimm-GAL4
UAS-Prosβ2, Prosβ6
Amino acid deprived
30 ℃
E
ns
ns
10000
8000
6000
4000
2000
0
Number of Sips Yeast
dimm-GAL4; tub-GAL80ts
UAS-Fbp1 RNAi

Figure S4. Related to Figure 4.

(**A-B**) No change in yeast intake was measured in protein deprived flies expressing RNAi against autophagy-related genes *Atg5* (A) or *Atg8* (B) in peptidergic neurons using *dimmed-Gal4* driver. n = 46-47 flies per group. (**C-D**) No change in yeast intake was measured in protein deprived flies expressing RNAi against *Prosα7* (C) or *Prosβ2* and *Prosβ6* dominant negatives (D) in peptidergic neurons using *dimmed-Gal4* driver. n = 47-48 flies per group. (**E**) No change in yeast intake was measured in protein deprived flies expressing RNAi against *Fbp1* in peptidergic neurons using *dimmed-Gal4* driver. n = 47-48 flies per group. ns $p \geq 0.05$, ** $p < 0.01$, Kruskal-Wallis ANOVA with Dunn's multiple comparisons test. Individual data points are single flies. Barplots are mean +/- standard error of the mean (SEM).

### Figure S5

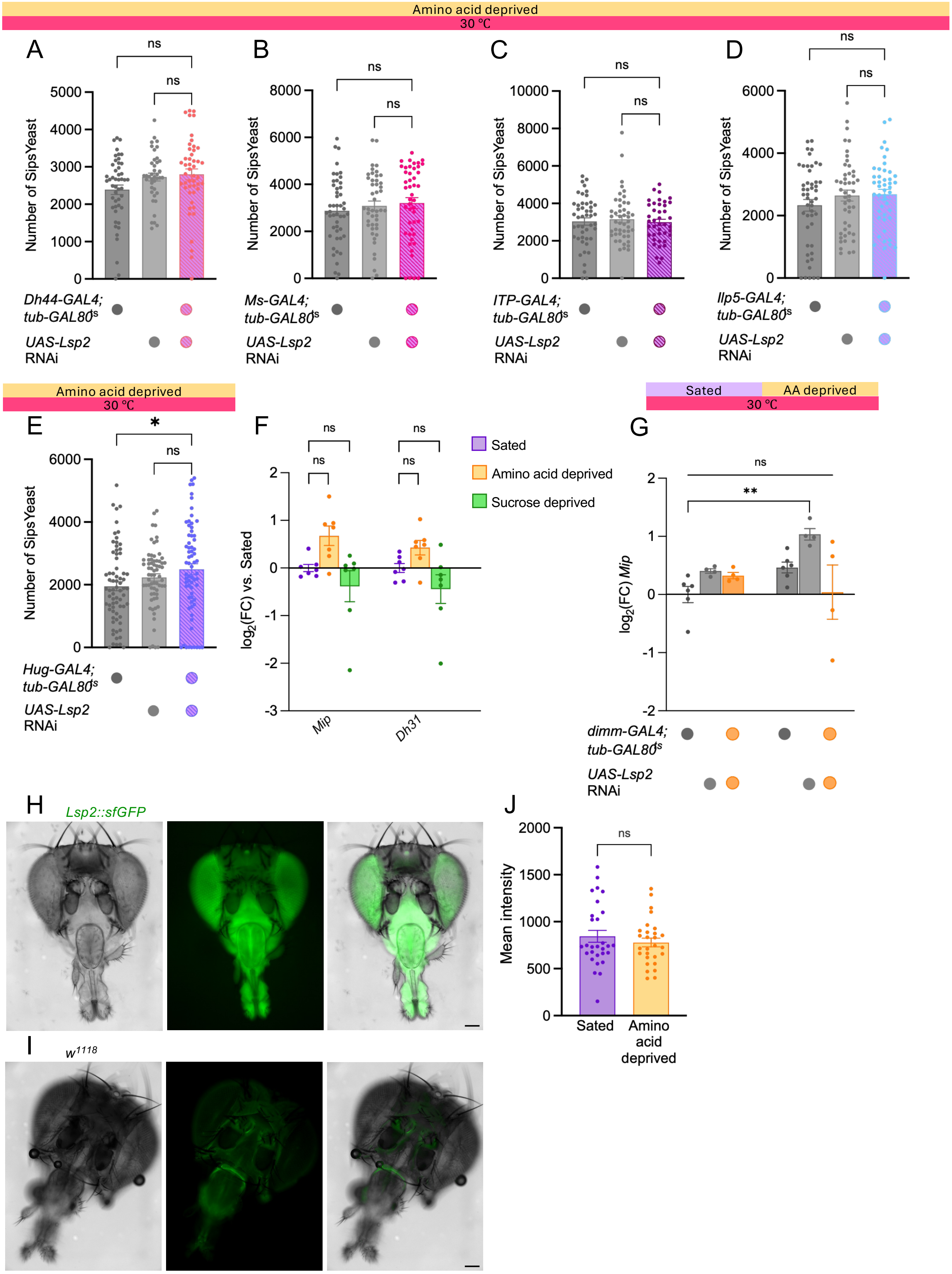

Amino acid deprived
30 ℃
A
B
C
D
ns
Number of SipsYeast
Dh44-GAL4; tub-GAL80ts
UAS-Lsp2 RNAi
Ms-GAL4; tub-GAL80ts
ITP-GAL4; tub-GAL80ts
Ilp5-GAL4; tub-GAL80ts
E
*
Hug-GAL4; tub-GAL80ts
F
Sated
Amino acid deprived
Sucrose deprived
log2(FC) vs. Sated
Mip
Dh31
G
Sated
AA deprived
**
log2(FC) Mip
dimm-GAL4; tub-GAL80ts
H
Lsp2::sfGFP
I
w1118
J
Mean intensity
Sated
Amino acid deprived

Figure S5. Related to Figure 5.

(**A-E**) No change in yeast intake was measured in protein deprived flies expressing *Lsp2* RNAi in *Dh44*+ (A), *Ms*+ (B), *ITP*+ (C), *Ilp5*+ (D) and *Hug*+ (E) neurons. n = 41-71 flies per group. (**F**) RT-qPCR showed no significant change in *Mip* transcript levels in the heads of amino acid or sucrose deprived flies, relative to sated flies. n = 6-7 groups of 25 flies each. (**G**) RT-qPCR showed no significant change in *Mip* transcript levels in the heads of sated or amino acid deprived flies with *Lsp2* RNAi expressed in *dimm-Gal4* neurons, relative to control flies. n = 4-6 groups of 25 flies each. (**H-I**) Representative brightfield (left), fluorescence (centre) and composite (right) images of fly heads expressing endogenously tagged Lsp2::sfGFP (H) or from *wild type* (*w1118*) control (I). (**J**) Quantification of mean fluorescence intensity in proboscises from sated and amino acid deprived flies. n = 28-29 flies per group. ns $p \geq 0.05$, * $p < 0.05$, ** $p < 0.01$, Kruskal-Wallis ANOVA with Dunn's multiple comparisons test. Individual data points are single flies, except in (F) and (G) where data points are groups of 25 flies. Barplots are mean +/- standard error of the mean (SEM).

### Figure S6

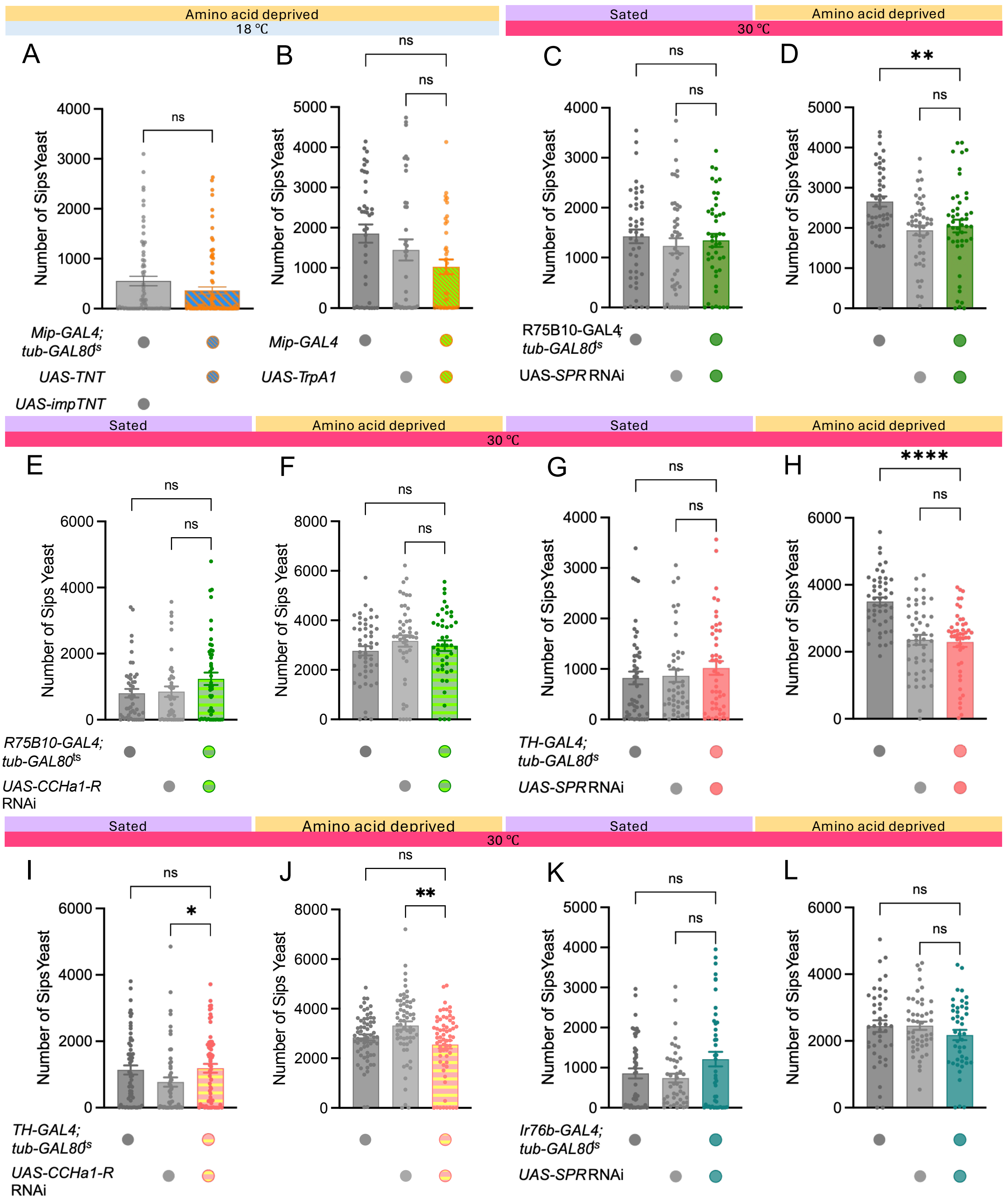

Amino acid deprived
18 ℃
Sated
Amino acid deprived
30 ℃
A
B
C
D
Number of Sips Yeast
ns
**
****
*
Mip-GAL4; tub-GAL80ts
UAS-TNT
UAS-impTNT
Mip-GAL4
UAS-TrpA1
R75B10-GAL4; tub-GAL80ts
UAS-SPR RNAi
E
F
G
H
UAS-CCHa1-R RNAi
TH-GAL4; tub-GAL80ts
I
J
K
L
Ir76b-GAL4; tub-GAL80ts

Figure S6. Related to Figure 6.

(**A**) Flies carrying *Mip-Gal4*, *tub-GAL80ts* and *UAS-TNT* kept at permissive temperature (18ºC) consume similar amounts of yeast as controls carrying *UAS-impTNT*. n = 74-77 flies per group. (**B**) Flies carrying *Mip-Gal4*, *tub-GAL80ts* and *UAS-Trpa1* kept at permissive temperature (18ºC) consume similar amounts of yeast as genetic controls. n = 38-40 flies per group. (**C-D**) No change in yeast intake was measured in sated (C) or amino acid deprived (D) flies expressing RNAi against *SPR* in FB-LAL neurons using *R75B10-Gal4* driver. n = 45-47 flies per group. (**E-F**) No change in yeast intake was measured in sated (E) or amino acid deprived (F) flies expressing RNAi against *CCHa1-R* in FB-LAL neurons using *R75B10-Gal4* driver. n = 42-48 flies per group. (**G-H**) No change in yeast intake was measured in sated (G) or amino acid deprived (H) flies expressing RNAi against *SPR* in dopaminergic neurons using *TH-Gal4* driver. n = 44-48 flies per group. (**I-J**) No change in yeast intake was measured in sated (I) or amino acid deprived (J) flies expressing RNAi against *CCHa1-R* in dopaminergic neurons using *TH-Gal4* driver. n = 55-64 flies per group. (**K-L**) No change in yeast intake was measured in sated (K) or amino acid deprived (L) flies expressing RNAi against *SPR* in amino acid sensing neurons using *Ir76b-Gal4* driver. n = 43-48 flies per group. ns $p \geq 0.05$, * $p < 0.05$, ** $p < 0.01$, Kruskal-Wallis ANOVA with Dunn's multiple comparisons test. Individual data points are single flies. Barplots are mean +/- standard error of the mean (SEM).
